# Deciphering spatial heterogeneity of patient-derived organoids of colorectal tumors for drug discovery

**DOI:** 10.64898/2026.02.10.704651

**Authors:** Maxim Norkin, Jonathan Bac, Louis McConnell, Christophe Cisarovsky, Marina Alexandre-Gaveta, Sylvie Andre, Simon Tanaka, Antonia Digklia, Sylke Hoehnel, Stephanie Tissot, Joerg Huelsken, Marianna Rapsomaniki, Raphael Gottardo, Krisztian Homicsko

**Author notes:** Equal contributors (Maxim Norkin, Jonathan Bac). Equal contributors (Raphael Gottardo, Krisztian Homicsko).

## Abstract

The spatial organization of tumors is critical for drug development, as spatial heterogeneity in cancer cells and the tumor microenvironment can present a significant barrier to effective therapies. Patient-derived tumor organoids (PDOs) offer promising platforms to study tumor biology and for drug development. Previous studies have demonstrated that PDOs can maintain the genomic/clonal heterogeneity of tumors, as well as expression heterogeneity. However, the spatial heterogeneity of PDOs and its correlation with the original tumors remains unclear.

We propose an integrated pipeline combining spatial transcriptomics and phenotypic analyses of PDOs to capture and track spatial heterogeneity. This platform integrates multiplexed organoid cultures with imaging-based spatial transcriptomics and analysis at single organoid resolution. We validate the platform by comparing results with non-spatial single-cell transcriptomic data and spatial transcriptomic analysis of donor tumors. The combination of *ex vivo* PDOs and spatial transcriptomics could open novel avenues for developing drugs.

## Introduction/Main

Patient-derived organoids (PDOs) have emerged as powerful preclinical models that recapitulate key features of the original tumor, including genetic alterations, histological architecture, and drug response profiles [1–3]. However, conventional bulk molecular profiling and non-spatial single-cell RNA sequencing (scRNA-seq) approaches fail to capture the spatial organization and morphological diversity inherent to organoid cultures [4,5]. This limitation hinders our ability to link cellular states with spatial architectures and to understand how spatial context influences tumor phenotypes.

Recent advances in spatial transcriptomics now enable high-resolution mapping of gene expression in space within intact tissues [6,7]. Technologies such as 10X Genomics Xenium provide subcellular resolution transcript detection across hundreds of genes, bridging the gap between whole-transcriptome scRNA-seq and spatially resolved molecular profiling [1].

When applied to organoid systems, spatial transcriptomics could elucidate how cellular programs are organized within individual organoids, how morphological features correspond to distinct molecular states, and how inter-organoid heterogeneity manifests at the spatial level [9,10].

Colorectal cancer (CRC) displays extensive heterogeneity across multiple scales—between patients, within individual tumors, and across metastatic sites—driven by diverse genetic, epigenetic, and microenvironmental influences [11,12]. This heterogeneity critically shapes tumor behavior, therapeutic response, and clinical outcome, yet remains insufficiently characterized at the cellular and spatial levels [2]. Notably, CRC was the first cancer type for which patient-derived organoids (PDOs) were established and remains one of the most well-characterized and widely used organoid model systems [14,15]. PDOs recapitulate the mutational landscape of original tumors ([3]). Despite their broad use in colorectal cancer research, most studies employing patient-derived organoids (PDOs) have relied on bulk transcriptomic, genetic, or functional readouts, which average signals across cells and obscure spatial organization of malignant programs([4], [5]). Studies comparing the PDOs’ and patient responses are still scarce and demonstrate only a limited prediction ([6]), most likely due to high levels of tumor and PDO heterogeneity. More recently, single-cell RNA sequencing has revealed extensive cellular heterogeneity and plasticity within CRC tumors and PDOs. Several malignant programs have been identified in PDOs such as goblet-like, enterocytes, proliferative transit-amplifying, crypt-like, inflammatory, EMT-like, metabolic and immune enriched ([7], [78], [73]). Furthermore, more extensive comparisons of tumor-PDO cell programs have been performed for several models highlighting a high level of malignant cell heterogeneity ([8], [34]). These approaches, however, dissociate cells from their native architecture, precluding analysis of spatial patterning and neighborhood relationships. Consequently, how malignant cell states are spatially organized within PDOs and how this organization compares to patient tumors, and whether spatial architecture contributes to functional heterogeneity and drug response remain largely unexplored.

Several tumor classification frameworks have been developed to improve understanding of colorectal cancer biology and guide therapeutic strategies. Among the most widely used are consensus molecular subtypes ([9]) and colorectal cancer intrinsic subtypes ([10]). CMS classification is derived from bulk tumor transcriptomes and therefore captures signals from malignant, immune, and stromal compartments, while CRIS is based exclusively on epithelial cell–intrinsic transcriptional programs and thus might be more relevant for organoid-based studies. CMS1 is characterized by hypermutated BRAF-mutant tumors, and high immune infiltration. CMS2 represents the canonical subtype with strong Wnt signaling and high proliferation levels, while CMS3 displays a metabolic phenotype and the highest prevalence of KRAS mutations. CMS4 is defined by a mesenchymal program and extensive stromal enrichment. Partial concordance exists between CMS and CRIS: for example, CRIS-C and CRIS-E largely mapping to CMS2, and CRIS-D capturing CMS4-like tumors. Despite its biological relevance, the clinical utility of CMS stratification has shown inconsistent results across clinical trials ([11], [79], [80]).

PDOs represent a powerful model for studying patient-specific drug responses; however, the contribution of the tumor microenvironment and its presence in ex-vivo cultures remains an open question. Several studies have demonstrated that cancer-associated fibroblasts (CAFs) can modulate PDO transcriptional profiles, although these effects are heterogeneous and not observed across all PDO-CAF combinations [[12] and [10]]. CAF co-culture has also been reported to partially influence responses to certain drug treatments, yet the extent of this effect remains incompletely understood [[10]]. Moreover, several models of PDO growth have been introduced: classical domes ([13]), constructed tumoroids ([14]) or larger cell aggregates ([15]). Each method offers its advantages and disadvantages: classical domes enable tracking organoid development from a single cell though lacking scalability, while tumoroids and cell aggregates provide size- and morphology-controlled structures compatible with high-throughput drug testing.

To date, only a limited number of studies have examined spatial heterogeneity in organoids, primarily relying on immunohistochemistry (IHC) to highlight a small set of markers (ttps://doi.org/10.3390/biology11091270, [75]). Imaging based spatial transcriptomic methods often focused on tissue development ([17], [81]). As a result, a comprehensive understanding of the spatial distribution of malignant cell states and programs within organoids, and the patterns related to matched tumors remain unresolved.

To overcome these limitations, we developed a spatially enabled multi-embedding platform that supports parallel spatial profiling of organoids across diverse experimental conditions, including developmental time courses, drug perturbations, and tumor–stromal co-cultures. By integrating Xenium spatial transcriptomics with whole-exome sequencing, whole-transcriptome single-cell RNA sequencing (scRNAseq, 10x Chromium), and clinical annotations, our platform enables systematic resolution of organoid heterogeneity across morphological, molecular, and functional dimensions. Using this approach, we generated a spatial transcriptomic atlas of 3,100 patient-derived organoids from 38 colorectal cancer organoid lines, alongside matched tumor tissue from 15 patients. Integrative analysis of spatial profiles from PDOs and tumors reveals shared malignant programs, delineates their spatial organization and heterogeneity, defines distinct organoid morphological classes, and links these features to drug treatment responses.

## Results

### A Single-Cell Spatial Atlas of Colorectal Cancer Organoids

In order to spatially profile CRC-derived PDOs, we generated a collection of 38 PDO lines based on established protocols (Figure 1A). In addition, when possible we also cultured cancer-associated fibroblasts (CAFs) from the same patients [18] (supp. Table 1). All CRC PDOs were early passages, passaged at least twice. The cohort of PDOs is representative of diverse clinical contexts of CRCs: 36% were derived from liver metastases, 37% from primary colon tumors, and 27% from primary rectal cancers. The majority, 70%, were from the left-side of the colon and 30% from the right side. Pathological analysis showed varied grades of differentiation with intermediate the most abundant (Supp. Table 1). To compare the genomic make-up of PDOs with known CRC genetics, we performed whole-exome sequencing of PDOs. This revealed the expected CRC driver mutations which included APC (80%), TP53 (60%), KRAS (50%), and BRAF (10%) (Figure 1B). These mutations were accompanied by characteristic chromosomal alterations: e.g. amplifications of CCND1, MYC, and EGFR (Supplementary Figure 1A). The transcriptional heterogeneity of CRC tumors has been previously defined by the Consensus Molecular Subtypes (CMS) and CRC intrinsic subtype (CRIS) classifications. Therefore, we next performed scRNAseq of the PDO lines and assigned CMS or CRIS signatures, as well as the microsatellite instability (MSI) status (Figure 1B) ([20]). As expected, PDOs with CMS3 and CRIS-A/E signatures were enriched in KRAS-mutant samples, while PDOs with predominant CMS2 and CRIS-C signatures were KRAS wild-type and TP53 mutant organoids. UMAP projection of the scRNA-seq data revealed predominantly patient-specific clustering of PDOs, with no clear segregation by KRAS/BRAF mutation status (Figure 1C and Figure 1E) or tumor stage (Figure 1D), indicating robust integration of gene expression profiles across organoids.

**Figure 1.**
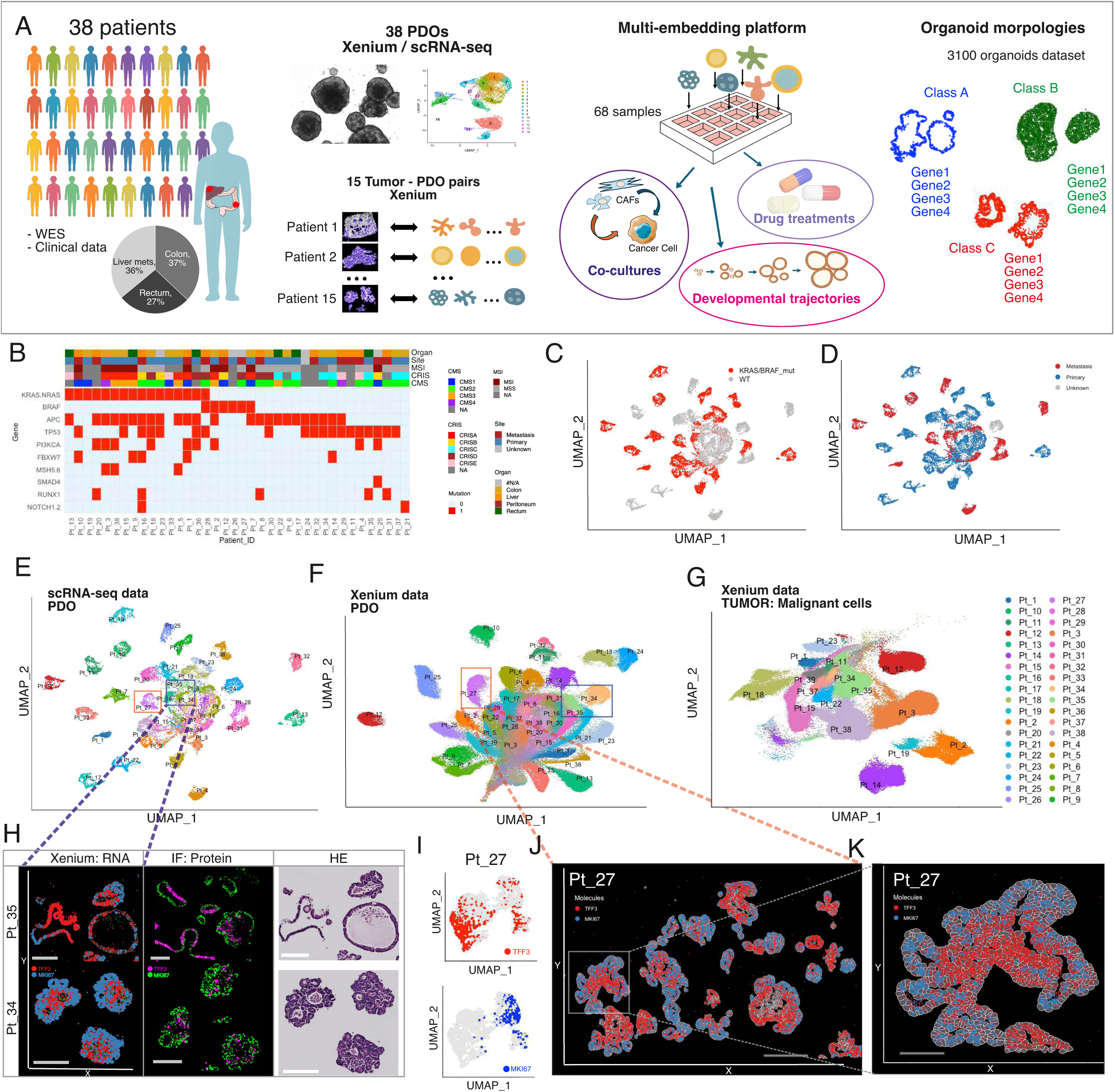
Non-spatial and spatial single-cell RNA analysis of patient-derived organoids from colorectal cancer patients. **A**. Schematic overview of the study. Left: Overview of biobank comprising 38 PDO lines with clinical annotations and multi-platform analysis including WES, scRNA-seq (Chromium), and Xenium spatial transcriptomics for 15 tumor-organoid pairs. Middle: Multi-embedding platform set-up description. Right: gene expression-based assessment of organoid morphologies. **B**. Clinical and molecular characteristics of the donors including anatomical site, MSI status, CRIS classification, and CMS subtype. (**C-E**) scRNAseq data based UMAP of PDOs color coded by mutation status, tumor site type and individual donors. (**F-G).** UMAP based on Xenium PDOs (F) and tumors (G) color coded by donor. **H**. Spatial plots showing the comparison of the *MKI67* / *TFF3* signals from Xenium (RNA), adjacent slides stained for the corresponding proteins and HE staining of the same region. Scale bars are 200 μm. **I**. UMAP based on scRNAseq data for donor Pt_27 showing *TFF3* and *MKI67* positive cells. **J-K**. Xenium spatial images demonstrating spatial organization of proliferative (*MKI67*) and differentiated (*TFF3*) states within individual organoids for patient 27, Pt_27. Scale bars are 200 um and 60 μm.

Because conventional scRNA-seq lacks spatial context, intrapatient and intra-organoid heterogeneity may be obscured. Therefore we performed imaging-based spatial transcriptomics on PDOs (Supplementary Figure 1B) with the Xenium platform (10X Genomics). To maximize detection sensitivity ([76]), we employed the pre-designed Immuno-Oncology panel comprising 380 genes (Supplementary Table 2). This analysis generated a spatially resolved atlas comprising around 550 ’000 cells across 2432 individual organoids (Figure 1F), with a median of 55 individual organoids per PDO line. On average, approximately 17,000 cells were profiled per PDO sample, with robust transcript detection (median of 90 genes and 430 transcripts per cell; Supplementary Figure 1C). Both scRNAseq (Figure 1E) and Xenium datasets exhibited substantial donor-to-donor variability. Xenium data recapitulated key transcriptional patterns observed in scRNA-seq (Supplementary Figure 1D). As PDOs are ex vivo models from tumors, we also profiled donor tumors from 15 of the 38 PDOs (Figure 1G). To illustrate the added value of Xenium’s spatial resolution, we highlight two PDO lines (Pt_34 and Pt_35), which exhibit spatially distinct expression patterns of MKI67, a marker of proliferation, and TFF3, a marker of goblet cell differentiation (Figure 1H). The spatial patterns of RNA expression closely mirrored the corresponding protein distributions (Figure 1H, middle panel), supporting the validity of the spatial transcriptomic approach. Another PDO line, Pt_27, shows pronounced intrapatient and intraorganoid spatial heterogeneity that is not captured by scRNA-seq. Although scRNA-seq identifies cells with differential expression of MKI67 and TFF3 (Figure 1I), spatial transcriptomics reveals a more nuanced organization at the level of individual organoids (Figure 1J–K). Within organoids from that same patient, we observed a continuum of cellular states ranging from highly proliferative (MKI67-high) to more differentiated (TFF3-high) cells, which were spatially segregated. Proliferative cells were preferentially localized to the outer layers—reminiscent of an invasive tumor front—whereas differentiated cells predominantly occupied inner regions. These spatially organized expression patterns are not detectable by scRNA-seq, highlighting the unique biological insights enabled by Xenium’s single-cell spatial resolution compared with bulk or non-spatial single-cell approaches.

### Single-cell spatial transcriptomic profiling of tumor-organoid pairs reveals shared patient-specific programs

To assess how faithfully organoids recapitulate malignant tumor cell states, we analyzed Xenium spatial transcriptomic data from 15 primary tumors matched to PDOs. Across these samples, we detected on average ∼55,000 cells per tumor (Figure 2A). The performance of the assay on tumor tissues and organoids was similar, with a median of 140 genes per cell on tumor samples compared to 90 genes per cell of PDOs. However, malignant cancer cells—comprising approximately 35% of all cells—showed higher transcriptomic complexity, with a median of 180 genes per cell, double that of PDOs.

**Figure 2.**
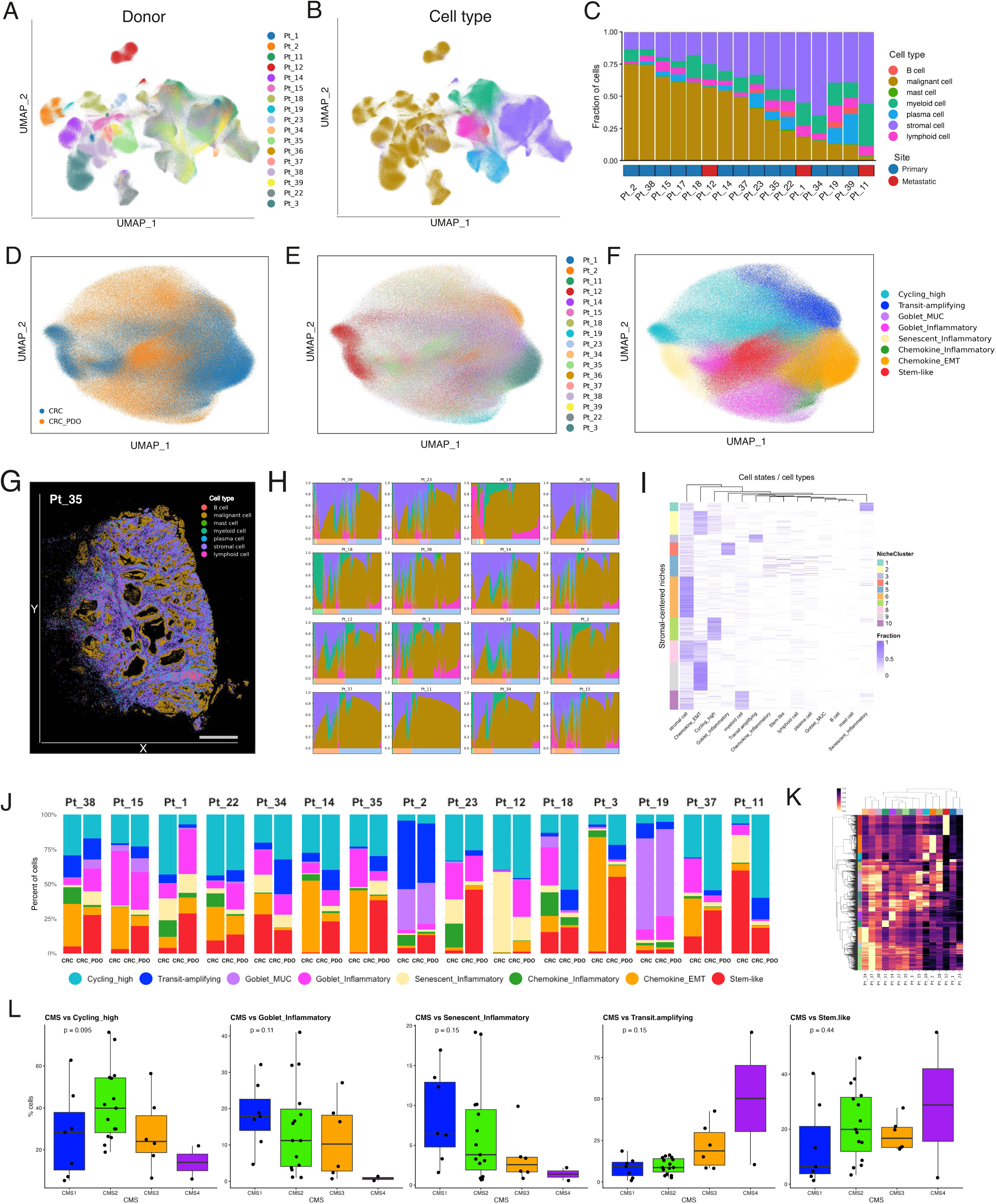
Comparison of donor tumor and PDO spatial profiles. **A-B**. UMAP integration of original tumor cells colored by patient identity and present cell types. **C**. Bar plots summarizing the frequency of annotated cell types for each donor’s tumor. **D-F**. UMAP of BBKNN-integrated Xenion single cells across tumor and PDO highlighted by tissue of origin, donor label, cell state programs identified with Leiden clustering. **G**. An example of a spatial image for the tumor (Pt_35 donor). Cell types are highlighted. The scale bar is 1 mm. **H**. Stromal cell-centered cell niche analysis based on the 15 μm radius. Cell niches composed of multiple cell types are present (niches containing only one cell type are filtered out). **I**. Niche compositions as a heatmap. **J**. Stacked bar plots comparing cellular program abundance between CRC tumors (left) and PDOs (right). **K**. Heatmap of cosine similarities between pseudo bulk tumors and corresponding PDOs using all overlapping genes. **L**. Fractions of BBKNNcell state programs between CMS subtypes calculated based on pseudo-bulked scRNA-seq expression data with CMScaller R package.

Cell-type labels were transferred from published single-cell RNA-seq data ([21]) using the Robust Cell Type Decomposition (RCTD) method ([22]) (Figure 2B). We could readily identify and quantify multiple cell types across the tumors: malignant cells, stromal cells (including fibroblasts and endothelial cells), plasma cells, mast and myeloid cells, lymphoid (T, NK cells) and B cells. Stromal and immune populations clustered by cell type across patients, whereas malignant cells clustered predominantly by patient of origin, consistent with strong inter-patient transcriptional differences (Figure 2B). Cell-type composition varied markedly within tumors and between patients, underscoring substantial intra-tumoral and inter-patient heterogeneity (Figure 2C, Figure 2G and Figure 2H).

To enable a direct comparison of malignant cells from tumors and PDOs, these cells were jointly classified into functional programs following BBKNN-based integration of the two modalities (see Methods) (Figure 2D). The resulting joint UMAP embedding revealed that malignant cells clustered predominantly by patient of origin rather than by sample type, with tumor- and organoid-derived cells intermingling within shared, patient-specific clusters (Figure 2E). These patterns indicate that, despite the absence of immune and stromal compartments in PDO cultures, malignant epithelial cell programs are somehow preserved and mirror those observed in primary tumors.

Leveraging the integrated space, we identified eight malignant cell programs shared between tumors and organoids, each defined by distinct marker gene expression profiles (Figure 2F, Supplementary. Figure 2A and Supplementary Table 3): Cycling_high (*MKI67*, *UBE2C*, *CDK1*), Transit-amplifying (*C1QBP*, *DMBT1*, *TUBA1B*), Stem_like (*RNF43*, *STAT6*), Goblet_MUC (*MUC5AC*, *REG4*, *ANPEP*), Goblet_Inflammatory (CEACAMs, *FOS*, *VEGFA*), Senescent_Inflammatory (*CDKN2B*, *CDKN2A*, *ANXA1*), Chemokine_EMT (*FN1*, *SPON2*, CXCLs, CCLs) and Chemokine_Inflammatory (*CXCL1*, *CXCL2*, *CXCL3*).

Malignant cells exist in specific neighborhoods, which could influence cancer cell development and their gene expression profiles (Figure 2G). Compositional neighborhood niche analyses centered on malignant cells revealed pronounced heterogeneity in the tumor microenvironment across patients (Figure 2H). Tumors from some patients exhibited highly heterogeneous niche compositions, characterized by variable immune and stromal infiltration (e.g., Pt_38, Pt_18, Pt_23, Pt_1), while others were characterized by more uniform microenvironments with limited non-malignant cell diversity (e.g. Pt_11, Pt_37, Pt_35). Overall, spatial organization analyses demonstrated a high degree of stromal fibroblast infiltration surrounding malignant cells across many tumors (Figure 2H).

Stratifying malignant cells by transcriptional programs within these spatial niches uncovered distinct microenvironmental associations. In particular, fibroblasts and myeloid cells were frequently co-localized with malignant cells, suggesting a supportive role in tumor growth and maintenance (Figure 2H). Notably, stromal cells—predominantly fibroblasts—were preferentially enriched in the neighborhoods of malignant cells in the Chemokine_EMT program (Figure 2I). These fibroblasts expressed high levels of *IL6*, *IL1B*, and *TGFB1*, consistent with activation of an IL1/TGFβ–CXCL signaling axis that induces the expression of CXC chemokine genes (*CXCL1*, *CXCL5*, *CXCL10*, *CXCL14*) in adjacent malignant cells.

The expression of CXC chemokines was further elevated in the Chemokine_Inflammatory malignant cell program, which localized to niches characterized by a combined enrichment of fibroblasts and myeloid cells, providing additional pro-inflammatory cytokine signals (Supplementary Figure 2B). Together, these findings indicate that local stromal and immune contexts are tightly coupled to specific malignant transcriptional programs, highlighting the role of spatially organized microenvironmental interactions in shaping tumor cell programs.

Comparing malignant cell programs between primary tumors and patient-derived organoids revealed overall concordance in the relative frequencies of cells assigned to each program (Figure 2J). Nonetheless, systematic differences in the abundance of specific malignant programs were observed between tumor- and organoid-derived cells. In particular, malignant cells in tumors exhibited higher proportions of Chemokine_EMT and inflammatory cell programs, consistent with the presence of substantial immune (myeloid, T, and B cells) and stromal (fibroblasts and endothelial cells) compartments in tumor tissue. As demonstrated by our niche analyses, these non-malignant populations are spatially co-localized with inflammatory and EMT-associated malignant programs but are absent from organoid monocultures, as expected.

Conversely, organoids exhibited an increased proportion of Cycling_high cells compared with tumors, likely reflecting selection for enhanced self-renewal and proliferative capacity under growth factor–rich culture conditions. Such quantitative differences between tumors and PDOs are therefore expected consequences of their distinct microenvironmental contexts. Despite these differences, assessment of global transcriptional concordance using cosine similarity across all genes revealed a preferential overall match between PDOs and their corresponding tumors (Figure 2K), suggesting that PDOs preserve the overall transcriptional features of their tumors of origin. CMS and CRIS molecular classifications are frequently used as descriptors of CRC heterogeneity; therefore we examined whether CMS and CRIS subtypes of colorectal cancers (Figure 1B) were associated with malignant transcriptional programs observed in PDOs (Figure 2L, Supplementary Figure 2E). As expected, Cycling_high cells were enriched in CMS2, while the Goblet_MUC program was enriched in CRIS-A subtype, consistent with the mucinous and differentiated epithelial nature of those tumors (Supplementary Figure 2F).

In summary, PDOs and tumors exhibit a high degree of malignant cell programs inheritance, with quantitative differences largely dictated by tumor microenvironmental influences and *ex vivo* culture conditions, underscoring the utility of PDOs for modeling patient-specific tumor spatial biology.

### Morphological diversity is associated with distinct molecular programs

Organoids exhibit pronounced heterogeneity in both morphology and transcriptional programs, even among organoids derived from the same donor. Moreover, these programs display a high degree of spatial organization within individual organoids (Figure 3A). Quantification of intra-patient program diversity using Shannon entropy revealed systematic associations with specific cell programs: Cycling_high programs were enriched in low-entropy organoids, whereas inflammatory-associated programs increased with higher entropy. This relationship was observed both at the individual organoid level and when aggregated at the donor level (Figure 3B, Supplementary Figure 3A), indicating that transcriptional heterogeneity is a reproducible feature across scales.

**Figure 3.**
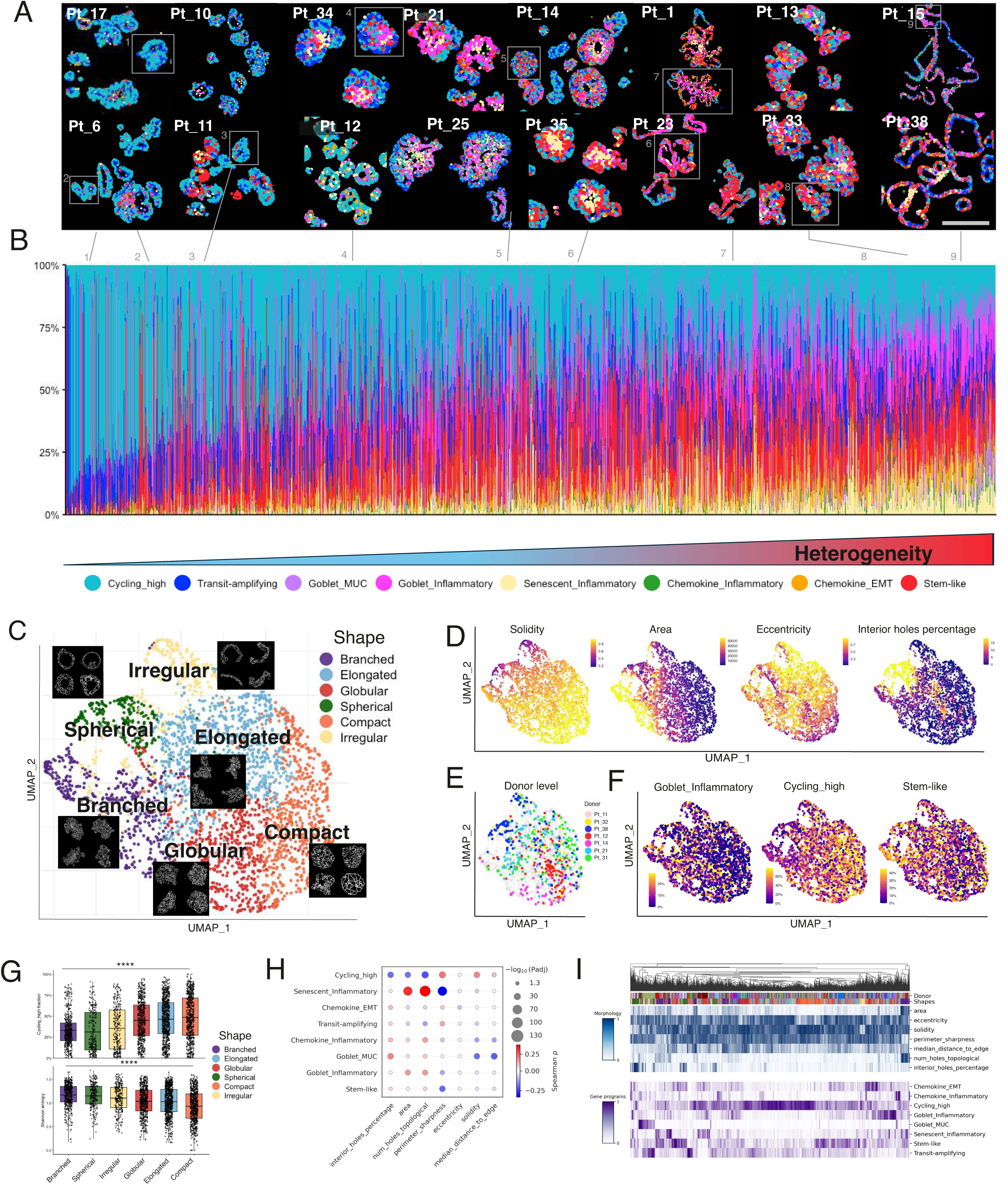
Molecular programs and morphological diversity of individual organoids. **A.** Spatial images of PDOs, cell state programs are highlighted. Organoids are organized by heterogeneity level (calculated Shannon entropy). The scale bar is ∼200 μm. **B**. Bar plots highlighting organoid program fractions and the link between selected organoids and their positioning in the bar plot is shown. Organoids are organized by the heterogeneity level (calculated Shannon entropy). **C**. UMAP projection of organoid shapes based on geometric features (e.g., area, eccentricity, solidity, interior holes percentage) colored by six morphological classes: branched, elongated, globular, spherical, compact and irregular with representative images for each class. **D-F**. Shape UMAP overlays colored by geometric features demonstrating quantitative morphological descriptors; showing distribution of selected donors; showing distribution of Cycling-high, and Goblet_inflammatory and Stem-like cell states across the organoid shapes. **G.** Box plots showing the distribution of the Cycling-high fraction and the organoid heterogeneity by all morphological classes. **H**. Dot plot highlighting the link between organoid features and cell state programs. **I**. Heatmap showing the distribution of the organoid features and cell state programs.

Given detected spatial variability, we next asked whether organoid morphology is linked to malignant cell programs. To address this, we constructed a comprehensive morphological atlas by applying k-means clustering to quantitative shape descriptors—including solidity, area, eccentricity, and the proportion of interior holes—across all 3100 organoids. This analysis classified individual organoids into six discrete morphological classes—compact, globular, branched, elongated, irregular, and spherical (Figure 3C)—which could be mapped using quantitative shape descriptors (Figure 3D). Morphological heterogeneity varied substantially both across donors (Supplementary Figure 3B) and among PDOs derived from the same patient (Figure 3E). For example, PDOs from Pt_38 displayed a broad spectrum of structures, including branched, hollow, and elongated morphologies, whereas PDOs from other patients, such as Pt_12 or Pt_14, were dominated by more homogeneous compact or globular forms.

Importantly, morphological classes showed distinct enrichments for malignant cell programs. For instance, the Goblet_Inflammatory program was preferentially enriched in branched and spherical organoids, whereas Cycling_high cells were most abundant in compact and globular structures (Figure 3F). These enrichments resulted in significant associations between specific programs, morphological classes, and levels of heterogeneity, exemplified by the Cycling_high program (Figure 3G). Joint clustering of morphological features and transcriptional programs recapitulated these associations and further revealed that combined morphology–program profiles were largely organoid-specific, with minimal clustering by donor (Figure 3H-I).

Together, these results indicate that organoid morphology and malignant cell programs are associated but not deterministically linked. Additional factors—such as internal organoid architecture and radial gene expression patterns—are likely to contribute to organoid formation and phenotypic diversity.

### Radial gene expression gradients define organoid architecture and cellular programs

Tumor development is a complex, multistep process that progresses from early lesions to large, spatially organized tumor masses. While early-stage lesions can vary in their balance between proliferation and dormancy, established tumors typically exhibit a conserved architectural pattern characterized by a differentiated, often hypoxic core surrounded by a proliferative outer front. We first assessed the presence of this organization using spatial expression of *MKI67* and *TFF3* in tumors, and observed that MKI67^+^ highly proliferative cells and TFF3^+^ differentiated cells showed spatial separation (Figure 4A).

**Figure 4.**
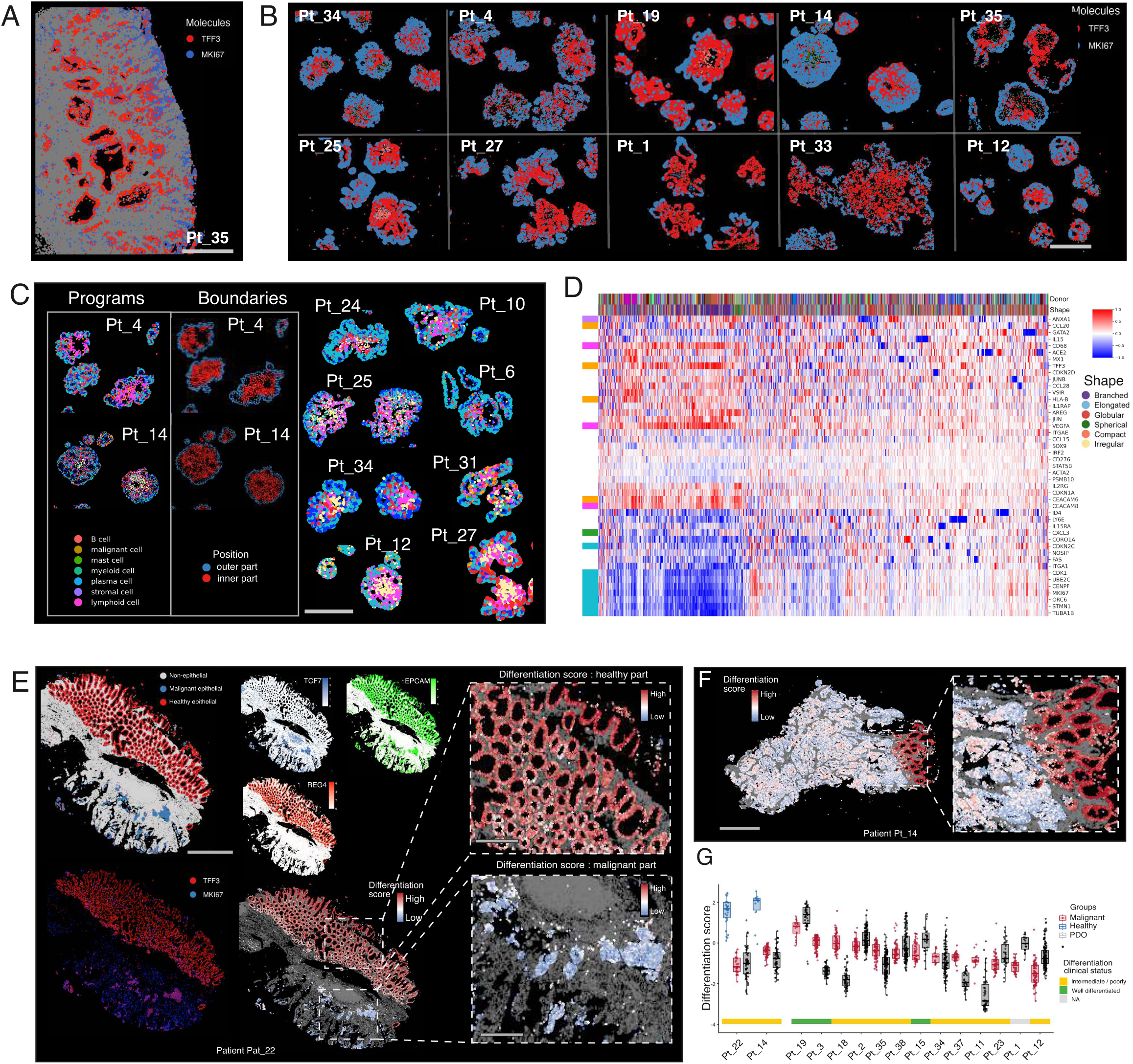
Organoid architecture reveals radial gene expression gradients. **A.** A representative image of the tumor showing spatial distribution of proliferative (*MKI67*) and differentiated (*TFF3*) markers, a tumor front. The scale bar is 1 mm. **B**. Representative organoid images showing spatial distribution of proliferative (*MKI67*) and differentiated (*TFF3*) markers. The scale bar is 200 μm. **C**. Representative organoid images showing: left = spatial distribution of the cell state programs; center - the boundaries definition (outer and inner organoid parts); right: more examples of the organized spatial distribution of the cell states. The scale bar is ∼200 μm. **D**. Heatmap showing top differentially expressed genes between outer and inner organoid layers and their affiliation with one of the cell states. **E-F**. Spatial images of the tumors (donor Pt_22 and Pt_14) consisting of healthy and malignant epithelial parts and their assessment with differentiation scoring. The scale bar is 1200 μm and 400 μm on the main and zoomed-in images on F, and 1 mm on G. **G**. **Left**: two donor examples that have healthy, malignant epithelial parts and PDOs. **Right**: Box plots highlighting the differentiation score of the tumors and PDOs for multiple donors, organized by decreasing median differentiation score. Horizontal bar annotations are showing the clinical status of those tumors.

We next asked whether patient-derived organoids recapitulate this higher-order spatial organization, beyond globally preserving malignant cell programs. Consistent with the tumor architecture, organoids from multiple donors exhibited a spatial separation of expression, with a TFF3^+^ core and an MKI67^+^ proliferative outer layer (Figure 4B).

To further investigate local organoid architecture at the transcriptional program level and confirm that this pattern extends beyond a small number of marker genes, we partitioned each organoid into inner and outer cellular layers and performed differential expression analyses between these regions (See Methods for details). In parallel, we spatially mapped malignant cell programs within individual organoids (Figure 4C). This analysis revealed a consistent spatial pattern: Cycling_high cells were preferentially localized to the outer regions of organoids, whereas inflammatory (e.g., *VEGFA*, *JUN*) and differentiated programs (e.g., *TFF3*, *CEACAM* family members, *ACE2*) were enriched in the organoid core (Figure 4C, Figure 4D). This behavior can also be observed in tumors (Supplementary Figure 4A).

Although this spatial organization was broadly conserved across organoid morphologies, the strength of the pattern varied by shape. Branched organoids exhibited the most pronounced radial organization, whereas spherical and irregular organoids showed weaker gradients (Figure 4D, Supplementary Figure 4B). These differences likely reflect the interplay between physical constraints, such as limited multilayered architecture, and culture-dependent factors including nutrient diffusion and growth factor availability, which together shape spatial zonation within organoids.

### Differentiation programs in tumors and PDOs

Building on the observed spatial distribution of proliferative and differentiation markers within organoids, we next asked whether clinically annotated tumor differentiation status could be inferred from Xenium gene expression profiles in primary tumors and their matched patient-derived organoids (PDOs). To address this, we analyzed specimens containing both malignant regions and adjacent normal epithelium (Figure 4E, Figure 4F). We computed a differentiation score for multiple tumor regions and corresponding organoids based on curated sets of differentiation- and proliferation-associated genes (See Methods). As expected, adjacent normal epithelial regions exhibited high differentiation scores, whereas malignant regions scored substantially lower (Figure 4E, Figure 4F). We similarly calculated differentiation scores for individual PDOs and compared the tumors of origin and PDOs to clinical annotation of differentiation (Figure 4G). Expectedly, PDOs derived from the matched tissues displayed low differentiation scores, consistent with their malignant nature (Pt_22, Pt_14). Tumor samples with the highest differentiation scores corresponded to clinically annotated “well-differentiated” tumors (Pt_19, Pt_3). While many organoids closely matched malignant tumor regions, substantial variability in differentiation scores was observed, potentially reflecting regional sampling differences within the tumors and high level of organoid heterogeneity. Indeed, we observed pronounced intra-patient heterogeneity across dozens of donors, with larger organoids tending to be more differentiated (Supplementary Figure 4C). The effect was most pronounced for branched and globular structures, which is consistent with the organoid maturation process and represents an additional source of biological complexity.

### Multi-embedding platform for higher throughput analysis of organoids

PDOs have been praised as potential platforms for *ex vivo* drug validation and screening. Drug screening using spatial transcriptomics of PDOs could be an ideal platform to profile spatially heterogeneous PDOs. However, the cost and time required to individually analyse organoids by imaging based spatial transcriptomics could be restrictive. To enable larger-scale, systematic spatial profiling of PDOs across experimental conditions, we developed a 3D-printed multi-embedding platform capable of accommodating 18–21 organoid samples per Xenium slide (Figure 5A, Figure 5B and Supplementary figure 5A). This approach would substantially reduce experimental costs while increasing throughput, facilitating time-course analyses, drug screening, and co-culture experiments. To validate the utility of our platform, we profiled organoid developmental trajectories by sampling organoids at six time points spanning 4 days to 5 weeks of growth (Figure 5C, upper panel). As expected, the number of cells per individual organoid increased progressively over time (Figure 5C, lower panel). We first confirmed our earlier observation that larger, more mature organoids exhibit higher levels of *TFF3* expression (Figure 5D). At the same time, malignant cell programs remained remarkably stable across developmental stages (Figure 5E). Indeed, only 5.3% (20/380) of genes showed significant temporal changes (Supplementary Figure 5B), including a gradual increase in differentiation markers (*TFF3*, *CDKN2D*) and a concomitant decrease in fetal-associated and metabolic genes (*LY6E, GLS2, SLC1A5, ACACA*).

**Figure 5.**
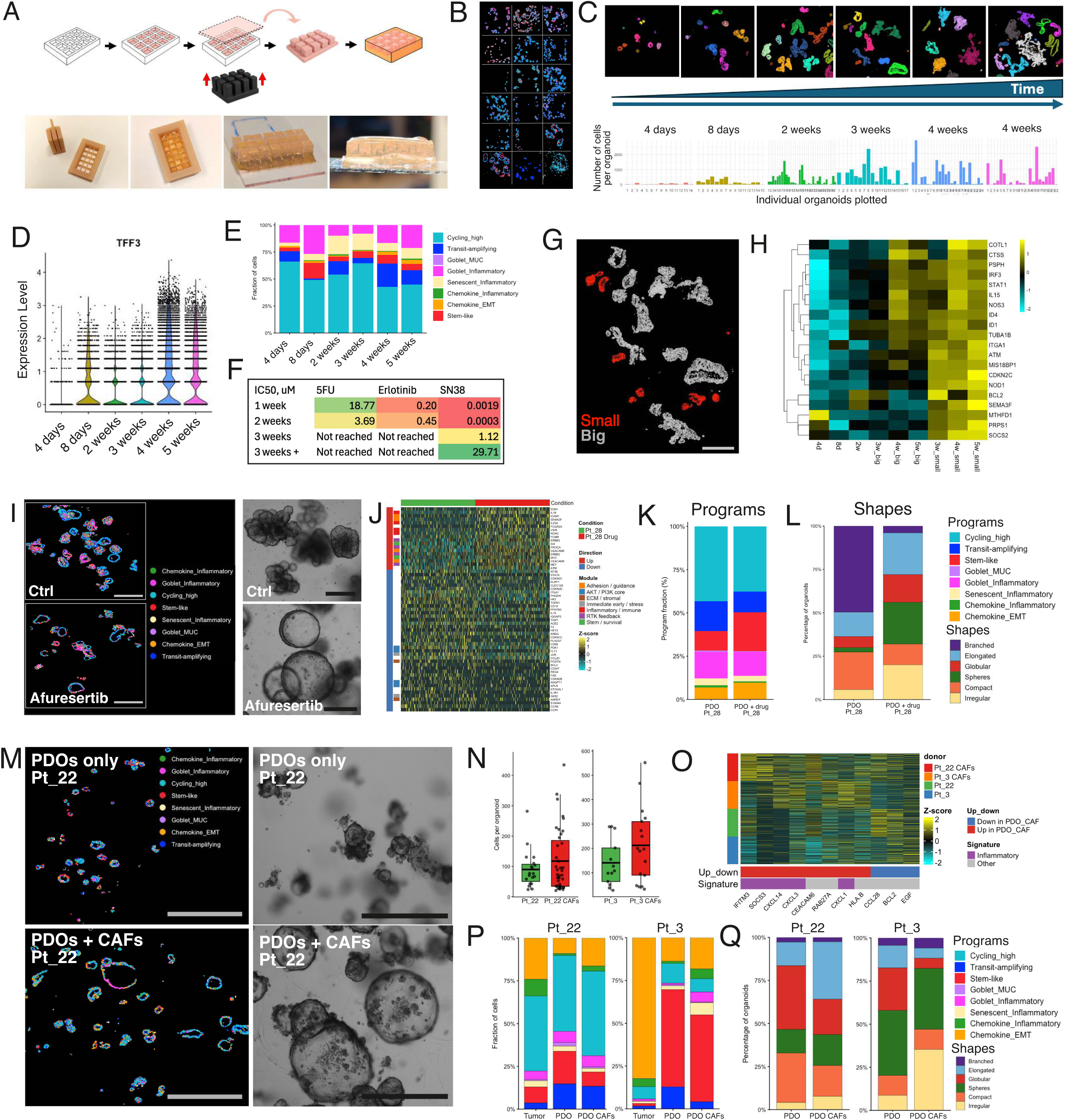
Multi-embedding platform enables high-throughput spatial profiling of organoid dynamics. **A**. Schematic representation and use and images of 3D-printed multi-embedding platform for parallel spatial transcriptomics. **B**. Representative Xenium field of view showing multiple embedded organoid samples. Cell programs are highlighted. Dimensions are 22×11 mm. **C**. Timeline of organoid developmental trajectory sampling from 4 days to 5 weeks with representative images showing size progression. Upper panel: representative spatial plots, organoid IDs are highlighted; bottom part: bar plots showing individual organoid sizes (cell numbers). **D**. Violin box plots showing cellular program composition across developmental stages. **E**. Cell program distributions across different stages of organoid development. **F**. IC50 results of several drug treatments across multiple stages of organoid development. Treatment duration - 3 days. **G**. Spatial Xenium plot showing identification of dormant organoids in late stages of organoid development. The scale bar size is 500 μm. **H**. Heatmap of developmental gene programs showing temporal dynamics of multiple gene programs through all developmental stages and between small and big organoids of later developmental stages. **I**. Spatial and BF images comparing PDOs alone versus drug treatment. Cell state programs are represented. The scale bar size is 250 μm for both Xenium spatial plot and BF image. **J**. Heatmap showing the most differentially expressed (DE) genes between control and drug treatment. **K-L**. Bar plot quantification of the cell state programs and organoid shapes by control and drug-treated conditions. **M**. Spatial and BF images comparing PDOs alone and PDO-CAF co-cultures for patient Pt_22. The bar size is 1200 μm for Xenium spatial plot and 500 μm for BF image. **N**. Bar plot quantification of organoid size between PDO and PDO + CAFs. **O**. Heatmap showing CAF-induced transcriptional changes in cancer cells for both donors. **P-Q**. Bar plot quantification of the cell state programs and organoid shapes by control and organoid from the CAF co-culture.

We next examined how the stage of organoid development might influence drug responses in order to define the ideal timing for drug testing. We tested established chemotherapies: 5-fluorouracil (5FU), active metabolite of irinotecan (SN38), as well as *EGFR* blockade by erlotinib. The treatment responses (IC50s) were largely consistent during the first two weeks of growth (Figure 5F). However larger and older (≥ 3 weeks) organoids displayed progressively increased resistance. Given the stable expression of organoid programs over time, this reduced sensitivity could be attributable to physical constraints on drug penetration in larger, more complex structures. Thus, although malignant cell-state programs remain stable over time, drug diffusion emerges as a critical determinant of therapeutic response, closely mirroring challenges encountered in the clinical setting.

Leveraging the spatial resolution of Xenium, we also identified a subset of small organoids that persisted over time (Figure 5G). These smaller organoids were transcriptionally distinct from larger counterparts, as well as the small organoids of early developmental stages (Figure 5H). We refer to this subset as dormant organoids, characterized by elevated expression of ID1 and ID4 transcription factors and interferon-stimulated genes (e.g., *STAT1*, *IFITM3*), consistent with a more quiescent transcriptional state despite prolonged culture and potentially increased treatment resistance despite their small size.

In previous work based on bulk RNA sequencing and phenotypic profiling of CRC organoids ([14]), the pan-AKT inhibitor afuresertib was identified as a candidate modulator of organoid morphology and gene expression. Building on these findings, we treated PDOs with afuresertib and profiled their responses using spatial transcriptomics. Consistent with prior observations, afuresertib treatment induced a pronounced morphological shift toward more spherical organoids, as detected by both brightfield (BF) imaging and Xenium analysis (Figure 5I). Differential expression analysis revealed upregulation of genes associated with cell adhesion, receptor tyrosine kinase feedback, and inflammatory signaling, mirroring changes previously observed by bulk RNA sequencing ([14]) (Figure 5J). While bulk RNA sequencing lacked the resolution for spatial programs, data from Xenium showed minimal change of cell-state programs following treatment (Figure 5K), in contrast to organoid morphology which shifted from predominantly branched to more spherical forms (Figure 5L).

Finally, to model tumor–stroma interactions, we performed spatial profiling of PDO–CAF co-cultures using the multi-embedding platform (Figure 5M). CAFs from co-culture were heterogeneous and segregated into two distinct clusters, which we called Type 1 and 2 (Supplementary Figure 5C). Type 1 CAFs expressed immune-suppressive iCAF and myofibroblast-like genes (e.g., *VEGFA*), whereas Type 2 CAFs were characterized by elevated expression of extracellular matrix (ECM) and antigen-presenting CAF-associated genes (Supplementary Figure 5D). The relative abundance of these CAF subtypes varied across patients (Supplementary Figure 5C). Co-culture of PDOs with CAFs resulted in larger organoids with increased cell numbers (Figure 5N), suggestive of growth support by CAFs. CAFs induced upregulation of chemokines in PDOs, including *CXCL1*, *CXCL3*, *CXCL14*, *CCL28* (Figure 5O), readily detectable from Xenium data. In addition, CAF co-culture altered malignant cell-state programs, with an increase in chemokine_EMT/inflammatory programs (Figure 5P). Organoid morphology was also affected, with a decrease in globular structures and an increase in elongated and irregular shapes (Figure 5Q). Together, the spatial profiling is compatible and informative on changes induced by CAFs on individual organoids.

## Discussion

We present a comprehensive spatial transcriptomic atlas of colorectal cancer patient-derived organoids, establishing fundamental principles of spatial organoid architecture and demonstrating the power of spatially resolved transcriptome profiling for the understanding of tumor heterogeneity and for future *ex vivo* drug testing. Through integration of Xenium spatial transcriptomics, single-cell RNA-sequencing, whole-exome sequencing and clinical data across 38 PDO lines and 15 matched tumor samples, we reveal that CRC organoids exhibit highly organized and heterogeneous spatial structures in which morphological diversity correlates with distinct molecular programs between patients but also within organoids of the same patient, reflecting inter- and intra-patient heterogeneity, respectively.

Spatial profiling enables deep characterization of developmental dynamics and tumor-stromal interactions. Furthermore, spatial programs, manifested as radial gene expression patterns in organoids are shared between PDOs and original donor tumors, underlying spatial reproducibility of expression heterogeneity. Proliferative, differentiated and inflammatory cells occupy distinct regions of PDOs which are specific for different groups of patients. This spatial pattern might reflect on fundamental biophysical constraints: cells in some organoids surface have preferential access to growth factors and nutrients in the culture medium, promoting proliferation, whereas core cells experience relative hypoxia and nutrient limitation, potentially triggering differentiation and metabolic adaptation [16–18]. The striking parallels between organoid radial zonation and tumor invasive front architecture suggest that organoids naturally recapitulate key aspects of *in vivo* tumor organization [19,20], validating their utility as models for spatial biology.

Our morphological atlas demonstrated a moderate association between morphology and shared gene programs across donors, revealing that morphology contains information complementary to gene expression alone. Organoid shape is not merely a descriptive feature but rather an informative phenotype partly linked to underlying molecular programs. [42,43]. These structure-function relationships mirror observations in tumor tissues, where architectural patterns correlate with clinical outcomes [48,49]. The ability to quantitatively profile organoid morphology through automated image analysis, combined with spatial transcriptomics, provides a scalable framework for high-throughput phenotypic screening [50,51]. Future studies could leverage this approach to identify morphology-modulating compounds or genetic perturbations, potentially revealing therapeutic targets that disrupt tumor architecture. Therefore, more efforts need to be undertaken to further improve organoid culture conditions, adapt to donor tumors. Comparison of matched tumor-organoid pairs revealed both preservation and divergence of cellular programs. Organoid cultures showed relative enrichment for stem-like and cycling populations compared to tumors [32,33], while chemokine-related cell states were diminished. This shift likely reflects selection for self-renewal capacity in culture and absence of microenvironmental constraints present *in vivo*.

The multi-embedding platform we developed addresses a practical bottleneck in spatial transcriptomics: the high cost and limited throughput of conventional workflows. By enabling parallel profiling of up to 21 conditions on a single Xenium slide, this platform facilitates time-course experiments, dose-response studies, and combinatorial screens that would otherwise be prohibitively expensive. In the future, we expect that even higher multiplexing could be achieved with further adaptation of 3D printing of cell containers. Our developmental trajectory analysis exemplifies this capability, revealing temporal gene programs that govern organoid maturation and identifying a subset of dormant organoids with distinct BMP signatures. These findings have implications for understanding tumor dormancy and therapy resistance, as quiescent cancer cells may evade treatment, seed recurrence [52–54] and should be targeted for efficient cancer therapies. Our detailed analysis of spatial cell architecture also lays the groundwork for future higher scale perturbation assays by optimizing the timing but also the critical endpoints parameters (shapes, expression, entropy) that should be taken into account when analysing organoid-based screens. Conversely, we would like to understand the importance of studying heterogenous and not homogenous organoid cultures. Efforts have been made to make the organoids more homogeneous and amenable for larger scale drug screens ([51]). Removing organoid heterogeneity might bias such homogenous assays and remove an important attribute of cancer biology.

The PDO-CAF co-culture experiments highlight the next level of complexity for organoid spatial research. CAFs induced inflammatory programs in cancer cells and altered organoid growth dynamics. While our current Xenium gene panel (380 genes) captured important immune and stromal markers, future studies using expanded panels or integrating Xenium with whole-transcriptome Visium data could provide even richer characterization of tumor-microenvironment crosstalk [55,56]. Such integrative approaches may identify targetable signaling axes mediating tumor-stromal interactions [55]. The PDO-CAF co-culture experiments showed a partial restoration of chemokine signatures in PDOs. Importantly, patient-specific program distributions were maintained in organoids, supporting their use as personalized disease models [59,60]. However, our findings also emphasize that organoid monocultures incompletely recapitulate the full tumor ecosystem—immune infiltration, stromal architecture, and vascular supply are absent [61,62]. Emerging organoid co-culture systems, including tumor-immune organoids and microfluidic models, promise to bridge this gap [63,64].

Several technical considerations merit discussion. First, the Xenium 380-gene immune oncology panel, while powerful for targeted profiling, captures only a fraction of the transcriptome. Genes relevant to metabolism, signaling, and other pathways may be underrepresented. Complementary whole-transcriptome spatial approaches (Xenium Prime, Visium HD) remain essential for comprehensive molecular characterization [57]. Second, organoid segmentation and morphological profiling rely on automated algorithms that may misclassify complex or crowded structures [65,66]. Manual curation and continuous algorithm refinement are necessary to ensure accuracy. Third, while our study encompasses 38 PDO lines—a substantial cohort—larger biobanks will be needed to fully capture the molecular diversity of CRC and enable robust biomarker discovery [4].

Looking forward, spatial transcriptomics holds immense promise for precision oncology. By profiling patient-derived organoids under diverse experimental conditions, we can systematically map genotype-phenotype relationships, predict therapeutic responses, and identify combination strategies [67,68]. The morphological and spatial features we describe may serve as biomarkers: for example, organoids with specific architectural patterns may predict sensitivity to targeted therapies or immunotherapies [69,70]. Integration of spatial profiling with high-throughput drug screens, CRISPR perturbation, and machine learning could accelerate discovery of predictive markers and therapeutic targets [71,72].

In conclusion, our work establishes spatial transcriptomics as a powerful tool for dissecting tumor heterogeneity in organoid models. We demonstrate that CRC organoids exhibit organized spatial structures shaped by conserved biological principles, that morphological diversity reflects distinct molecular programs, and that spatial profiling enables deep characterization of cellular dynamics and microenvironmental interactions. The datasets, analytical frameworks, and experimental platforms we present provide a foundation for future studies aimed at leveraging spatial biology to advance precision medicine in colorectal cancer and beyond.

## Methods

### Human tumor sample collection

Colorectal samples were obtained from the Biobank of the Department of Clinical Oncology of the Centre Hospitalier Universitaire Vaudois (CHUV). The protocol for sample collection and organoid generation was approved by the local ethics committee (CER-VD) study ID CHUV_DO_CTE_TRP_0001. All patients consented for sample collection and organoid generation.

### Patient derived organoid and CAFs generation

Tumor samples were washed in phosphate-buffered saline (PBS) and mechanically minced into small fragments using sterile blades. Subsequently, tumor fragments were enzymatically incubated in a digestion buffer containing collagenase II at 2.5 mg/ml (Sigma-Aldrich; C6885-1G), collagenase IV at 2.5 mg/ml (Sigma-Aldrich; C5138-1G), and DNase I at 0.5 mg/ml (Roche; #11284932001) in advanced DMEM-F12 media supplemented with Y-27632, ROCK-I kinase inhibitor (10uM) for 40 minutes at 37°C on a gently rotating platform to enhance enzymatic activity. Following digestion, the tumor suspension was filtered through a 100 µm cell strainer. Fetal bovine serum (FBS) was immediately added to the collected fraction to neutralize enzymatic activity. The samples were centrifuged at 320 × g for 5 minutes, after which the supernatant was discarded. Cell pellets were washed once in with advanced DMEM-F12 media and centrifuged at 320 × g for 5 minutes. Cells were resuspended in ENR media (advanced DMEM-F12, B27 1x, N2 1x, HEPES 10 mM, Glutamax 1x, Primocin 1x, EGF (50 ng/ml), Rspo (titred, in-house production), Noggin (titred, in-house production), mixed with matrigel (Corning, 356231) and seeded in 48-well TC treated plates. After 15 minutes incubation at 37°C, matrigel domes were covered with 300 uL of ENR media.

CAFs isolation followed the same digestion protocol and obtained cell suspension was seeded in 6-well TC plate and 3 ml of the CAF media was added: DMEM supplemented with 10% FBS, 1x penicillin/streptomycin, 1 ng/mL of basic fibroblast growth factor and 5 ng/mL of insulin.

### Co-culture experiments and organoid treatments

Organoids were trypsinized to single cells or small cell clusters and mixed with CAFs in 1:1 ratio. Around 50 000 cells were seeded in a 48-well TC treated plate and grown for 2 weeks before harvesting both CAFs and PDOs.

PDO treatments were done at two week’s time points after splitting for the drug experiments in Figure 5I-K and Supp. Figure 3B-C. The drugs were applied for 3 days before the readout. Afuresertib concentration was 10 uM in Figure 5I-K experiment.

Drug treatments in developmental trajectories experiments were performed for 3 days and the corresponding organoids from each stage were used from the same passage as aliquotes.

### Whole-exome sequencing

Organoids were grown in 6 well TC plates, collected, trypsinized into single cells, and resuspended in 200 µL of PBS (200 000 - 500 000 cells). Genomic DNA was isolated from the corresponding samples using the NucleoSpin Blood Mini Kit (Macherey–Nagel, REF: 740951.50) according to the manufacturer’s instructions. Whole-exome capture was performed using the xGen Exome Research Panel v2 (IDT, REF: 10005152), together with the xGen Hybridization and Wash Kit (IDT, REF: 1080577) and xGen Universal Blockers-TS (IDT, REF: 1075474). Library preparation was carried out using the xGen DNA Library Prep EZ Kit (IDT, REF: 10009863).

Amplified libraries were sequenced on a NovaSeq SP platform using paired-end 150-bp reads, achieving an average sequencing depth of 84x (approximately 20 million reads per sample). Raw FASTQ files were trimmed using Trim Galore (v0.6.6) with default parameters (adapter sequence: 5′-AGATCGGAAGAGC-3′). Reads were aligned to the Homo sapiens GRCh38 primary assembly reference genome using BWA-MEM (v0.7.17). Resulting BAM files were generated and processed using SAMtools (v1.10).

Duplicate reads were marked, and read groups were added using Picard (v2.20.8). Base quality score recalibration was performed with GATK (v4.3.0.0) using known variant sites from the NCBI database (common_all_20180418.vcf.gz), followed by application of recalibration scores. Variant calling was conducted using GATK HaplotypeCaller, and resulting VCF files were consolidated, with SNPs and INDELs selection and filtered according to GATK best practices. Variant annotation was performed using snpEff (GRCh38.86). Final variant tables containing all high-confidence SNPs and INDELs were generated using the GATK VariantsToTable function. Clinically relevant variants were identified by cross-referencing with the ClinVar database of pathogenic variants and the OncoKB database, focusing on the 500 most frequently mutated genes in colorectal cancer.

### Single-cell RNA-sequencing

Organoids were trypsinized and strained through a 40um strainer. Obtained single cell suspension was further processed with 10X Genomics 3’ Cell Multiplexing with CMOs (CellPlex, 1000261) kit according to the manufacturer protocol. In brief, cells were washed with PBS and centrifuged at 400g for 5 minutes at 4°C. Cells were counted and stained with CellPlex oligos and passed through several rounds of washes (400g for 5 minutes at 4°C). Cells were counted again and mixed in the proportions according to the manufacturer protocol. Cells were applied on Chromium Next GEM Chip G and processed for cell encapsulation and library preparation on Chromium Controller. Libraries and CMOs were amplified and sequenced at the depth of 800 mio reads per 30 000 cells for the library and 200 mio reads per 30 000 cells CMOs using NovaSeq or SPOCK sequencing machines at 150 nt PE configuration.

### Single-cell RNA-sequencing data processing and analysis

Obtained fastq files were processed using cellranger (version 7.1.0) according to the manufacturer 10x 3’ Multiplex pipeline using GRCh38 human reference assembly with min-assignment-confidence parameter equal to 0.6. Cells with more than 20% of MT genes were filtered out. For the UMAP generations and clustering a lognorm transformation of the data was used (Seurat_5.3.0, Dimplot function).

Infercnv (R package) was used to predict gene amplifications and deletions based on single cell RNA sequencing data.

### Multi-embedding platform 3D-printing

The CAD models were designed using Onshape (https://www.onshape.com/) (freely licensed for academic, non-profit use) and are publicly accessible (https://cad.onshape.com/documents/01e00334364d2028b9f273e6/w/3edd98f50286789d95a2d101/e/000fe6326d9008364abdb770?renderMode=0&uiState=68fe62a53b5ef48e6a3fafe

6). The models were 3D printed using a Dentsply Sirona Primeprint printer and Post-Processing Unit (PPU) (https://www.dentsplysirona.com/en-us/discover/discover-by-brand/primeprint.html) with Dentsply Sirona Primeprint Model material (https://www.dentsplysirona.com/en-us/discover/discover-by-brand/primeprint/indications-and-materials.html) and standard settings.

The platform consists of a multi-pocket mold (15x, 18x, 21x samples) with open-bottom wells that matches Xenium slide dimensions (21×10 mm) and a matching plunger that ejects the gel cubes containing embedded organoid from the mold.

### Spatial transcriptomics data generation

Organoids were grown approximately 2 weeks or until their size reached 200-500 um. Organoids were collected and fixed in 4% PFA overnight at +4 degrees. Fixed organoids were sedimented and washed with PBT buffer (0.1% (vol/vol) Tween in PBS). Organoids were resuspended in 50-100 ul of Histogel (HG-4000-012) preheated to 55-60°C using 200 ul tip cut at the edge (to increase the nozzle size), the drops were placed on the cap of 10 sm dish and placed in the fridge (+4°C) for 5-10 minutes for solidification. Generated per-patient drops were processed for paraffin embedding using histokinette machine and embedded in paraffin blocks. Sections (6 µm) from 8–10 donor samples derived from separate paraffin blocks were mounted onto Xenium slides and processed according to the manufacturer’s protocol.

For multi-embedding platform–based processing, 18–21 samples were collected and prepared upfront (stored in PBST buffer following overnight PFA fixation) and processed simultaneously. A multi-pocket mold was secured onto a Superfrost slide using Scotch tape applied along its two shorter sides. Each sample was prepared individually: organoids were sedimented and the supernatant was removed. 50-100 ul of preheated Histogel was added to a single sample, gently mixed using 200 ul tips cutted at their edge, and transferred into one pocket of the mold. The slide was gently tapped several times to allow organoids to sediment within the mold well, after which it was placed on a pre-cooled metal rack (4-8°C) for 10–20 seconds to solidify the gel.

All 18–21 samples were processed sequentially in this manner. Once all mold pockets were loaded, additional preheated Histogel (500–1000 µL) was carefully added dropwise on top of the wells to link them together and form a single contiguous block. A plunger was then used to release the entire structure from the mold by pressing into the pockets. Subsequently, the spaces between the resulting columns were slowly perfused with an additional volume of preheated Histogel (∼500 µL), applied dropwise to avoid melting of the pre-formed columns. Finally, the complete rectangular structure was incubated in 70% ethanol for 48 h prior to paraffin embedding ( the step is critical to prevent shrinkage and cracking of the block). The embedded structure was sectioned at 6 µm and mounted onto Xenium slides for downstream processing.

### Xenium Data Preprocessing and Quality Control

The raw Xenium output was processed using the snakemake pipeline at https://github.com/bdsc-tds/xenium_analysis_pipeline/tree/c36425b14e9fd638f0eeb8430f72449a974e6b2e, for cell segmentation and ProSeg 2.0.0 software ([27]). Following segmentation, a quality control (QC) filter was applied to the cell-feature matrix. Cells were retained for downstream analysis only if they met the criteria of having a minimum of 20 total transcript counts and expressing at least 5 unique genes. This step ensures the removal of low-quality or poorly segmented cells from the dataset. Additionally, spatial regions of poor quality were cropped out based on a significant drop of genecounts (Supp. Figure 1C).

### Data analysis

#### Cell Type Identification and Compositional Data Analysis

To annotate cell types, we employed Robust Cell Type Decomposition (RCTD) ([22]). This method leverages a curated external single-cell RNA sequencing dataset as reference ([22]) to assign cell type labels to cells in our Xenium samples.

ProSeg segmentation showed the most accurate results when comparing the cell type assignment by RCTD to a manually curated tumor cell type classification (Supp. figure 2C). We compared several reference datasets for the cell type annotation and obtained similar results in terms of cell type abundances (Supp. figure 2D). RCTD annotation using several references highlighted the presence of multiple cell types across the tumors: malignant cells, stromal cells (including fibroblasts and endothelial cells), plasma cells, mast and myeloid cells, lymphoid (T, NK cells) and B cells (Figure 2B).

For the analysis of cellular neighborhoods, we counted cell type labels within a 15µm radius around each individual cell across all colorectal cancer (CRC) tumor samples, and divided by the total number of cells. For computational efficiency, the resulting matrix of proportions was then quantized into 1000 clusters using KMeans from scikit-learn, before hierarchical clustering using scipy’s linkage with Jensen-Shannon distance and ‘average’ method. The resulting tree was then cut to obtain 10 clusters.

#### Data Integration and Identification of Gene Programs

To create a unified view and compare cellular states across the different experimental conditions, we integrated data from both CRC tumor tissues and the tumor-derived organoids. The integration was performed on log-normalized ProSeg expected counts. After reducing dimensionality to the first 50 principal components, we utilized the Batch Balanced K-Nearest Neighbors (BBKNN) algorithm ([28]), specifying the sample of origin as the variable to integrate over.

Following integration, we used the Leiden algorithm at resolutions 0.5,0.6,0.7,0.8 to identify distinct malignant cell clusters present across all samples. After manual review and choice of 0.8 as the best resolution, we identified the characteristic gene programs for each cluster by differential expression analysis using a t-test, comparing the gene expression of cells within a cluster to all other cells. The top differentially expressed genes were used to define the gene signature for each program. The scanpy package was used to perform all the above described analyses ([29]).

#### Segmentation and Definition of Organoid Units

To analyze the spatial organization of cells within tumor-derived organoids, we developed a graph-based segmentation approach. First, a spatial graph was constructed where each node represents a cell. An edge was drawn between two nodes if their corresponding cell boundaries were touching or overlapping in the physical space.

We then identified the connected components within this graph, with each component representing a putative organoid. To refine these initial organoid definitions, a further two-step curation process was implemented. Leiden clustering was performed on the graph at resolution 1.0, and the results were used to split bigger components into single, coherent organoid units. Each component was manually reviewed and split using lasso selection instead if Leiden was not satisfactory. Components that were excessively small (<20 cells) were discarded.

#### Analysis of organoid cellular layers

To assign cells within a given organoid to layers, we used the geopandas and shapely library to identify the outer boundary of each organoid. Cells touching the boundary were defined as the 0th layer. Cells touching 0th layer cells were defined as 1st layer, and so on. Cells not touching the last layer were defined as “interior”. For each organoid, differential expression was performed between cells in layers 0 and 1 vs all others using scanpy.tl.rank_genes_group’s default method (t-test). To define top genes per shape, we took the top and bottom 10 genes per log fold-change after subsetting to genes with adjusted p-values <0.05 and median percent of expressing cells across organoids >40%.

#### Pseudobulk comparison of organoids and tumors

We computed pseudobulk profiles of individual PDOs, Xenium tumors and Chromium tumors by summing their counts before log-normalization. We then assessed the similarity of PDOs with their patient-matched Xenium and Chromium tumors using cosine similarity. We additionally used pseudobulk profiles of organoids to score CMS/ CRIS and MSI signatures using CMScaller R package (v. 2.0.1, [30]).

### Morphological Analysis

After organoid segmentation, we computed a series of morphological features for each organoid, namely solidity, area, eccentricity, perimeter, and empty space.

We generated masks by joining the ProSeg segmentations for each organoid unit rasterizing the underlying geometries of each cell segmentation. In order to standardize the input for downstream analysis, all masks are scaled to a fixed square window size. Because there is a very large size range in organoids, we observed that many metrics are heavily influenced by the absolute size of the organoid (major/minor axis length, perimeter, etc.). We found that generating masks to fill the middle 90% of the mask window enables better representation of the relative shape of the organoids.

To quantitatively evaluate the morphological properties of the organoid units, we calculated seven morphological metrics from scikit-image ([31]) derived from the binary masks, namely area, perimeter sharpness, eccentricity, solidity, median distance to edge, number of holes and percentage of interior holes. Area is defined as the pixel sum of the rescaled organoid unit, and perimeter sharpness is defined as the sum of all boundary pixels of the organoid unit, serving as an indication of boundary complexity. Structural elongation was characterized by the major and minor axis lengths of an ellipse with equivalent normalized second central moments and the proportion between these two metrics (eccentricity). To measure the amount of empty space within the organoid, we computed the extent (area of minimal bounding box / area of organoid) and the solidity (area of minimal convex bounding region / area of organoid).

## Data availability

Processed data of single-cell RNAseq, spatial transcriptomics and whole exome sequencing will be made available through GEO. For access to raw data, reasonable requests should be addressed to the corresponding authors.

## Code availability

The code to reproduce the analysis can be found at https://github.com/bdsc-tds/Norkin2026.

## Competing financial interest

The authors declare no financial interest.

## Funding

K.H. received funding for this project from Innosuisse and the ISREC Foundation. R.G. has received consulting income (payments made to the Lausanne University Hospital) from Takeda, Arcellx, Sanofi, Owkin, and declares ownership in Ozette Technologies. K.H received research funding from Bristol Myers Squibb, ROCHE/Genentech, Merck Sharp & Dohme, Tolremo AG and Boehringer Ingelheim. K.H. and R.G. have received research funding from Owkin and 10x Genomics.C.C received funding from the Swiss National Science Foundation (SPARK grant).

## Acknowledgements

We thank P. Dessen (EPFL, Lausanne) for growth factor production, Veronique Noguet (CHUV, Lausanne) for organoid mounting to the Xenium slides. We thank various facilities for their help in the various parts of the project: GTF UNIL core facility for the sequencing, AGORA facilities HF FBM UNIL for histology, BET EPFL facility for the microscopy.

## Author contributions

M.N., K.H. and R.G. conceived the project. M.N. and K.H. designed the research and M.N. performed the experiments. M.N., J.B. and L.M. established the bioinformatics pipelines, and performed the computational analysis, with contributions from R.G., K.H., M.R. M.N. and J.B. generated the figures. M.A-G., S.A. and K.H. have processed the Xenium slides. S.Tanaka have designed the multiembedding platform with the inputs from M.N. S.Tissot have performed IF experiments and help with the interpretation of the results. C.C. and A.D. contributed to sample acquisition and clinical data collection. C.C., S.H., J.H., M.R., R.G., K.H. provided financial support for the study. M.N., K.H., R.G. prepared the manuscript, with the scientific inputs from J.B., M.R., L.M., J.H., S.H.

## Notes

### Competing Interest Statement

The authors have declared no competing interest.

### Summary of Updates

1. Exchange signs for Maxim Norkin in the author list. (should be co-correspondence) 2. Figure 5 (current) should be Figure 2. 3. Add figure names on top/before each figure. 4. Add Figure annotations/captions. 5. Supp figures link doesn't work

## References

[1] Janesick A, et al. High resolution mapping of the tumor microenvironment using integrated single-cell, spatial and in situ analysis. Nat Commun. 2023;14(1):8353.

[2] Dienstmann R, et al. Consensus molecular subtypes and the evolution of precision medicine in colorectal cancer. Nat Rev Cancer. 2017;17(2):79–92.

[3] Kuilman T, et al. Preserved genetic diversity in organoids cultured from biopsies of human colorectal cancer metastases. Proc Natl Acad Sci U S A. 2015;112(43):13308–13311.

[4] van de Wetering M, et al. Prospective derivation of a living organoid biobank of colorectal cancer patients. Cell. 2015;161(4):933–945.

[5] Boj SF, et al. Organoid models of human and mouse ductal pancreatic cancer. Cell. 2015;160(1-2):324–338.

[6] Ooft SN, et al. Patient-derived organoids can predict response to chemotherapy in metastatic colorectal cancer patients. Sci Transl Med. 2019;11(513):eaay2574.

[7] Fujii M, et al. A colorectal tumor organoid library demonstrates progressive loss of niche factor requirements during tumorigenesis. Cell Stem Cell. 2016;18(6):827–838.

[9] Guinney J, et al. The consensus molecular subtypes of colorectal cancer. Nat Med. 2015;21(11):1350–1356.

[10] Isella C, et al. Selective analysis of cancer-cell intrinsic transcriptional traits defines novel clinically relevant subtypes of colorectal cancer. Nat Commun. 2017;8:15107.

[11] Stintzing S, et al. Consensus molecular subgroups (CMS) of colorectal cancer (CRC) and first-line efficacy of FOLFIRI plus cetuximab or bevacizumab in the FIRE3 (AIO KRK-0306) trial. Ann Oncol. 2019;30(11):1796–1803.

[13] Sato T, et al. Single Lgr5 stem cells build crypt-villus structures in vitro without a mesenchymal niche. Nature. 2009;459(7244):262–265.

[14] Pleguezuelos-Manzano C, et al. Mutational signature in colorectal cancer caused by genotoxic pks+ E. coli. Nature. 2020;580(7802):269–273.

[15] Boehnke K, et al. Assay establishment and validation of a high-throughput screening platform for three-dimensional patient-derived colon cancer organoid cultures. J Biomol Screen. 2016;21(9):931–941.

[16] Sasaki N, et al. Reg4+ deep crypt secretory cells function as epithelial niche for Lgr5+ stem cells in colon. Proc Natl Acad Sci U S A. 2016;113(37):E5399–E5407.

[17] Yvernault BC, et al. Mapping mesenchymal diversity in the developing human intestine and organoids. bioRxiv. 2025. doi: 10.1101/2025.07.22.665939.

[18] Sato T, et al. Long-term expansion of epithelial organoids from human colon, adenoma, adenocarcinoma, and Barrett’s epithelium. Gastroenterology. 2011;141(5):1762–1772.

[20] Sveen A, et al. Colorectal cancer consensus molecular subtypes translated to preclinical models uncover potentially targetable cancer cell dependencies. Clin Cancer Res. 2018;24(4):794–806.

[21] Qi L, et al. Integrative single-cell analysis of human colorectal cancer reveals patient stratification with distinct immune evasion mechanisms. Nat Cancer. 2024;5(9):1409–1426.

[22] Cable DM, et al. Robust decomposition of cell type mixtures in spatial transcriptomics. Nat Biotechnol. 2022;40(4):517–526.

[23] Jones DC, et al. Cell simulation as cell segmentation. Nat Methods. 2025;22(1):83–91.

[24] Chen Z, et al. Spatially resolved transcriptomics reveals genes associated with the vulnerability of middle temporal gyrus in Alzheimer disease. Acta Neuropathol Commun. 2022;10(1):188.

[25] Williams CG, et al. An introduction to spatial transcriptomics for biomedical research. Genome Med. 2022;14(1):68.

[26] van de Wetering M, et al. Prospective derivation of a living organoid biobank of colorectal cancer patients. Cell. 2015;161(4):933–945.

[27] Polański K, et al. BBKNN: fast batch alignment of single cell transcriptomes. Bioinformatics. 2020;36(3):964–965.

[28] Wolf FA, et al. SCANPY: large-scale single-cell gene expression data analysis. Genome Biol. 2018;19(1):15.

[29] Eide PW, et al. CMScaller: an R package for consensus molecular subtyping of colorectal cancer pre-clinical models. Sci Rep. 2017;7(1):16618.

[30] van der Walt S, et al. scikit-image: image processing in Python. PeerJ. 2014;2:e453.

[31] Broguiere N, et al. Dissection and reconstruction of the colorectal cancer tumor microenvironment. bioRxiv. 2023. doi: 10.1101/2023.11.01.565169.

[32] Drost J, Clevers H. Organoids in cancer research. Nat Rev Cancer. 2018;18(7):407–418.

[33] Vlachogiannis G, et al. Patient-derived organoids model treatment response of metastatic gastrointestinal cancers. Science. 2018;359(6378):920–926.

[34] Ganesh K, et al. A rectal cancer organoid platform to study individual responses to chemoradiation. Nat Med. 2019;25(10):1607–1614.

[35] Rao A, et al. Exploring tissue architecture using spatial transcriptomics. Nature. 2021;596(7871):211–220.

[36] Moses L, Pachter L. Museum of spatial transcriptomics. Nat Methods. 2022;19(5):534–546.

[37] Marx V. Method of the Year: spatially resolved transcriptomics. Nat Methods. 2021;18(1):9–14.

[38] Williams CG, et al. An introduction to spatial transcriptomics for biomedical research. Genome Med. 2022;14(1):68.

[39] Melo Ferreira R, et al. Integration of spatial and single-cell transcriptomics localizes epithelial cell-immune cross-talk in kidney injury. JCI Insight. 2021;6(12):e147703.

[40] Russell AJC, et al. Slide-tags enables single-nucleus barcoding for multimodal spatial genomics. Nature. 2024;625(7995):101–109.

[41] Nagle PW, et al. Lack of DNA mismatch repair combined with an inflammatory microenvironment drives adaptive resistance to immune checkpoint blockade. Nat Commun. 2023;14(1):3432.

[42] Hofer M, Lutolf MP. Engineering organoids. Nat Rev Mater. 2021;6(5):402–420.

[43] Tuveson D, Clevers H. Cancer modeling meets human organoid technology. Science. 2019;364(6444):952–955.

[44] Bronsert P, et al. Cancer cell invasion and EMT marker expression: a three-dimensional study of the human cancer-host interface. J Pathol. 2014;234(3):410–422.

[45] Hao Y, et al. Integrated analysis of multimodal single-cell data. Cell. 2021;184(13):3573–3587.

[46] Pastushenko I, et al. Identification of the tumour transition states occurring during EMT. Nature. 2018;556(7702):463–468.

[47] Nieto MA, et al. EMT: 2016. Cell. 2016;166(21):21–45.

[48] Lugli A, et al. Recommendations for reporting tumor budding in colorectal cancer based on the International Tumor Budding Consensus Conference (ITBCC) 2016. Mod Pathol. 2017;30(9):1299–1311.

[49] Dawson H, et al. Tumor budding/T-cell infiltrates in colorectal cancer: proposal for a novel combined score. Histopathology. 2020;76(4):572–580.

[50] Caicedo JC, et al. Data-analysis strategies for image-based cell profiling. Nat Methods. 2017;14(9):849–863.

[51] Bougen-Zhukov NM, et al. A suite of organoid models that recapitulate colorectal cancer liver metastasis enables drug response profiling. J Exp Clin Cancer Res. 2023;42(1):131.

[52] Recasens A, Munoz L. Targeting cancer cell dormancy. Trends Pharmacol Sci. 2019;40(2):128–141.

[53] Sosa MS, et al. Mechanisms of disseminated cancer cell dormancy: an awakening field. Nat Rev Cancer. 2014;14(9):611–622.

[54] Endo H, Inoue M. Dormancy in cancer. Cancer Sci. 2019;110(2):474–480.

[55] Sahai E, et al. A framework for advancing our understanding of cancer-associated fibroblasts. Nat Rev Cancer. 2020;20(3):174–186.

[56] Biffi G, Tuveson DA. Diversity and biology of cancer-associated fibroblasts. Physiol Rev. 2021;101(1):147–176.

[57] Rao A, et al. Exploring tissue architecture using spatial transcriptomics. Nature. 2021;596(7871):211–220.

[58] Moses L, Pachter L. Museum of spatial transcriptomics. Nat Methods. 2022;19(5):534–546.

[59] Ganesh K, et al. A rectal cancer organoid platform to study individual responses to chemoradiation. Nat Med. 2019;25(10):1607–1614.

[60] Vlachogiannis G, et al. Patient-derived organoids model treatment response of metastatic gastrointestinal cancers. Science. 2018;359(6378):920–926.

[61] Drost J, Clevers H. Organoids in cancer research. Nat Rev Cancer. 2018;18(7):407–418.

[62] Clevers H. Modeling development and disease with organoids. Cell. 2016;165(7):1586–1597.

[63] Dijkstra KK, et al. Generation of tumor-reactive T cells by co-culture of peripheral blood lymphocytes and tumor organoids. Cell. 2018;174(6):1586–1598.

[64] Neal JT, et al. Organoid modeling of the tumor immune microenvironment. Cell. 2018;175(7):1972–1988.

[65] Stringer C, et al. Cellpose: a generalist algorithm for cellular segmentation. Nat Methods. 2021;18(1):100–106.

[66] Greenwald NF, et al. Whole-cell segmentation of tissue images with human-level performance using large-scale data annotation and deep learning. Nat Biotechnol. 2022;40(4):555–565.

[67] Yao Y, et al. Patient-derived organoids predict chemoradiation responses of locally advanced rectal cancer. Cell Stem Cell. 2020;26(1):17–26.

[68] Tiriac H, et al. Organoid profiling identifies common responders to chemotherapy in pancreatic cancer. Cancer Discov. 2018;8(9):1112–1129.

[69] Schutte M, et al. Molecular dissection of colorectal cancer in pre-clinical models identifies biomarkers predicting sensitivity to EGFR inhibitors. Nat Commun. 2017;8:14262.

[70] Pauli C, et al. Personalized in vitro and in vivo cancer models to guide precision medicine. Cancer Discov. 2017;7(5):462–477.

[71] Ringel T, et al. Genome-scale CRISPR screening in human intestinal organoids identifies drivers of TGF-β resistance. Cell Stem Cell. 2020;26(3):431–440.

[72] Dempster JM, et al. Chronos: a cell population dynamics model of CRISPR experiments that improves inference of gene fitness effects. Genome Biol. 2021;22(1):343.

[73] Marteau V. et al., Single-cell integration and multi-modal profiling reveals phenotypes and spatial organization of neutrophils in colorectal cancer, Cancer Cell, 2026 Jan 12;44(1):

[74] Ganesh K. et al., A rectal cancer organoid platform to study individual responses to chemoradiation, Nat Med. 2019 Oct;25(10):1607–1614.

[75] Ma H. Multiscale Analysis of Cellular Composition and Morphology in Intact Cerebral Organoids, Biology 2022, 11(9), 1270

[76] Mariia Bilous, et al, From Transcripts to Cells: Dissecting Sensitivity, Signal Contamination, and Specificity in Xenium Spatial Transcriptomics, bioRxiv. 2025. doi: 10.1101/2025.04.23.649965]

[77] Jyoti Rao, Unified Generation of Regionalized Neural Organoids from Single-Lumen Neuroepithelium, bioRxiv. 2025. doi: 10.1101/2025.11.18.689013]

[78] Pelka K, et al. Spatially organized multicellular immune hubs in human colorectal cancer. Cell. 2021;184(18):4734–4752.e20.

[79] Okita A, et al. Consensus molecular subtypes classification of colorectal cancer as a predictive factor for chemotherapeutic efficacy against metastatic colorectal cancer. Oncotarget. 2018;9(27):18698–18711.

[80] Buikhuisen JY, et al. Exploring and modelling colon cancer inter-tumour heterogeneity: opportunities and challenges. Oncogenesis. 2020;9(7):66.

[81] Kozlowski D, et al. Genetically Engineered Brain Organoids Recapitulate Spatial and Developmental States of Glioblastoma Progression. Adv Sci. 2025;12(7):e2407763.

