## Supplementary figures and images for "Deciphering spatial heterogeneity of patient-derived organoids of colorectal tumors for drug discovery"

### Supplementary table 1

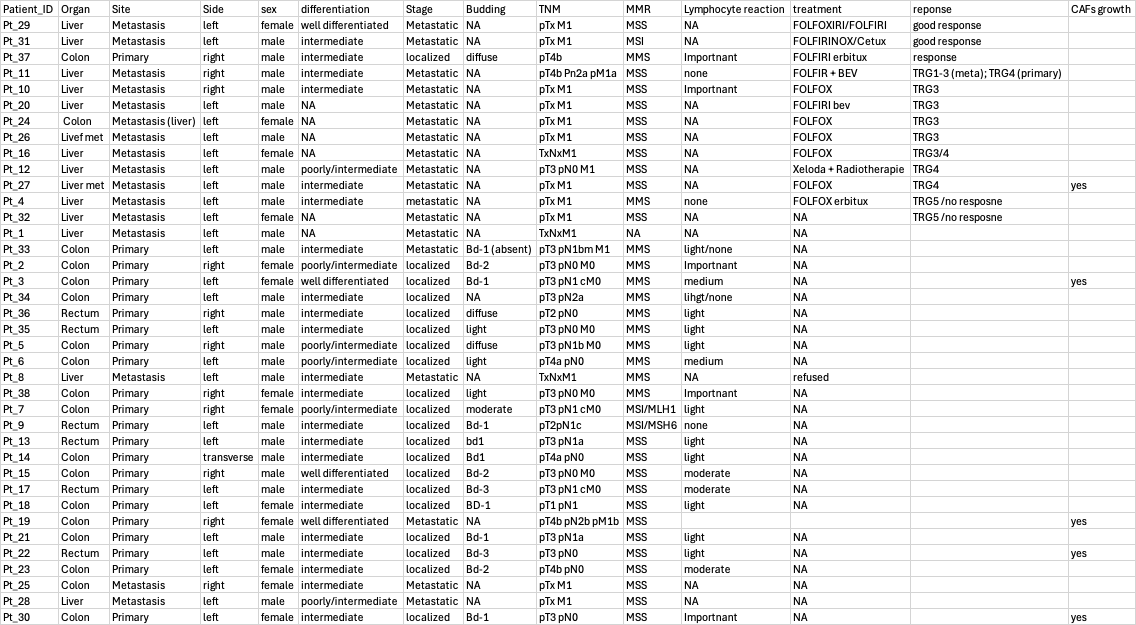
