## Supplementary Figures 1 to 5 for "Deciphering spatial heterogeneity of patient-derived organoids of colorectal tumors for drug discovery"

### Supplementary Figure 1

A

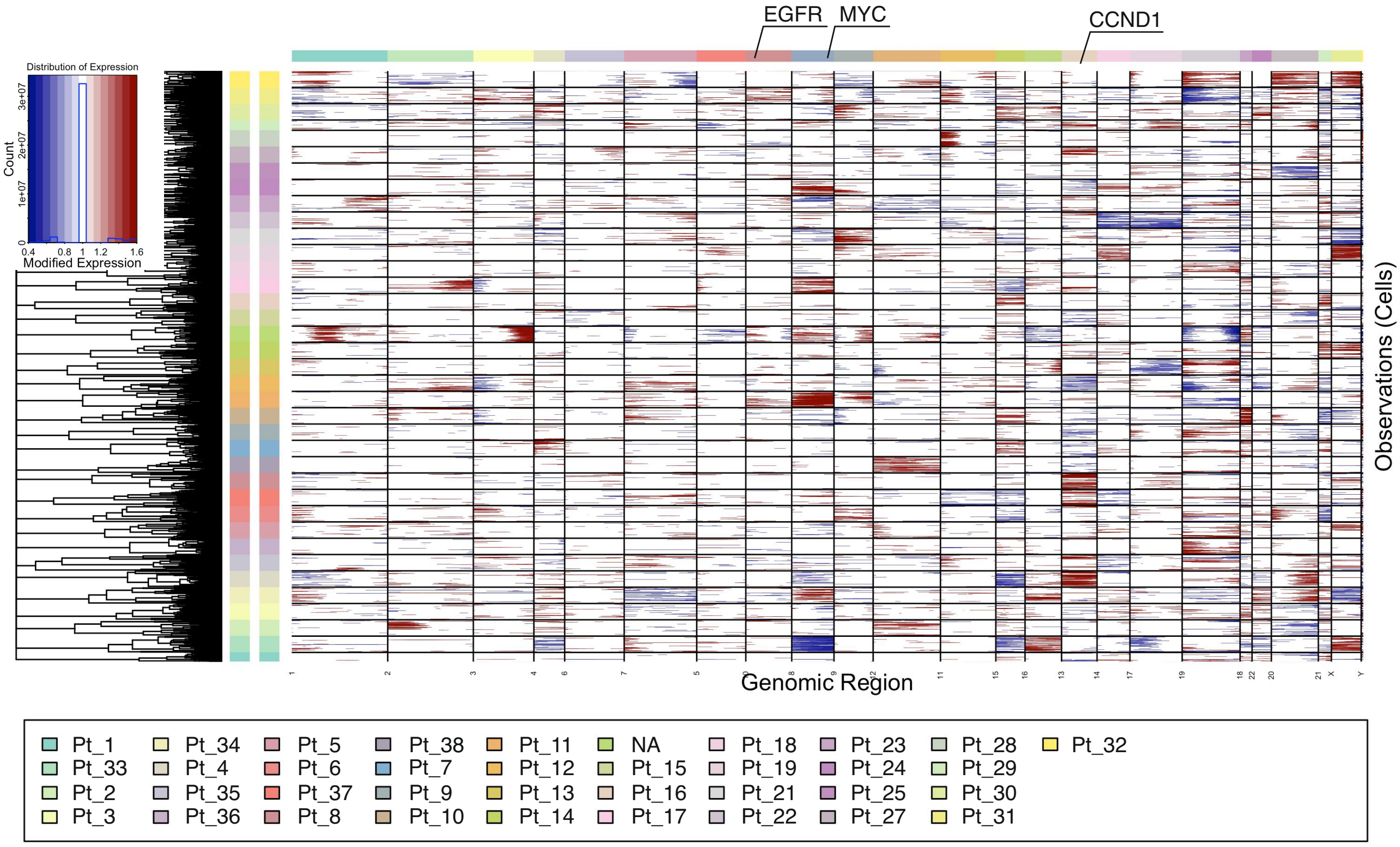


B

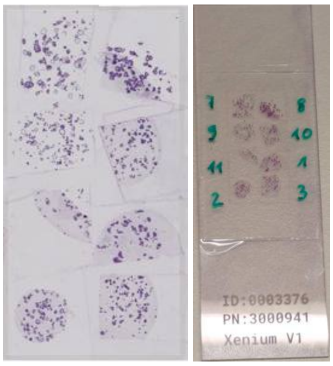


C

Pre-processing strategy:
removing the low count regions; cell filtering min_n_counts = 20

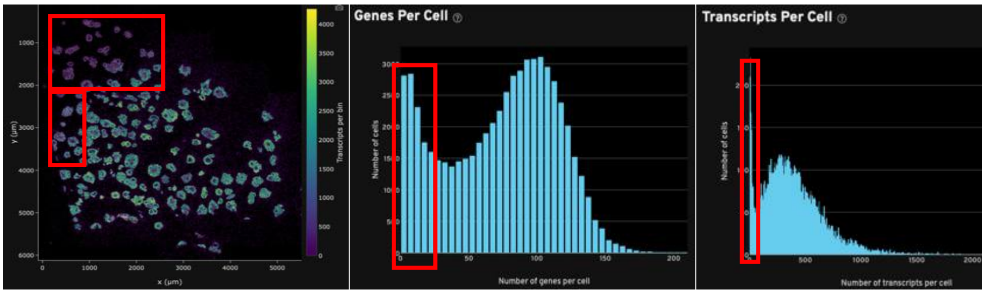


D

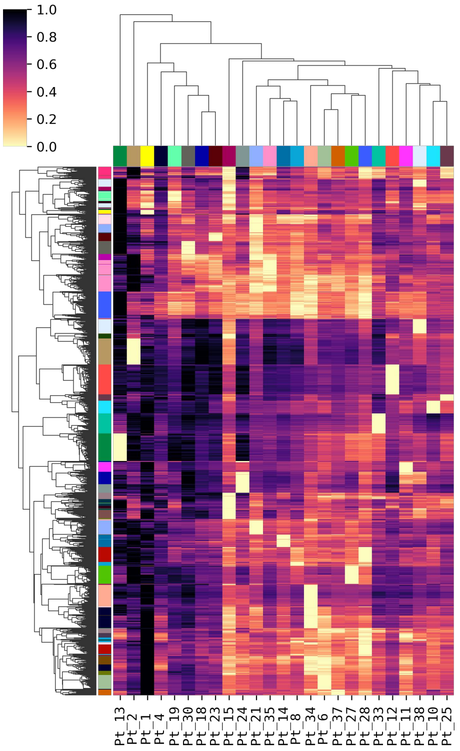

### Supplementary Figure 2

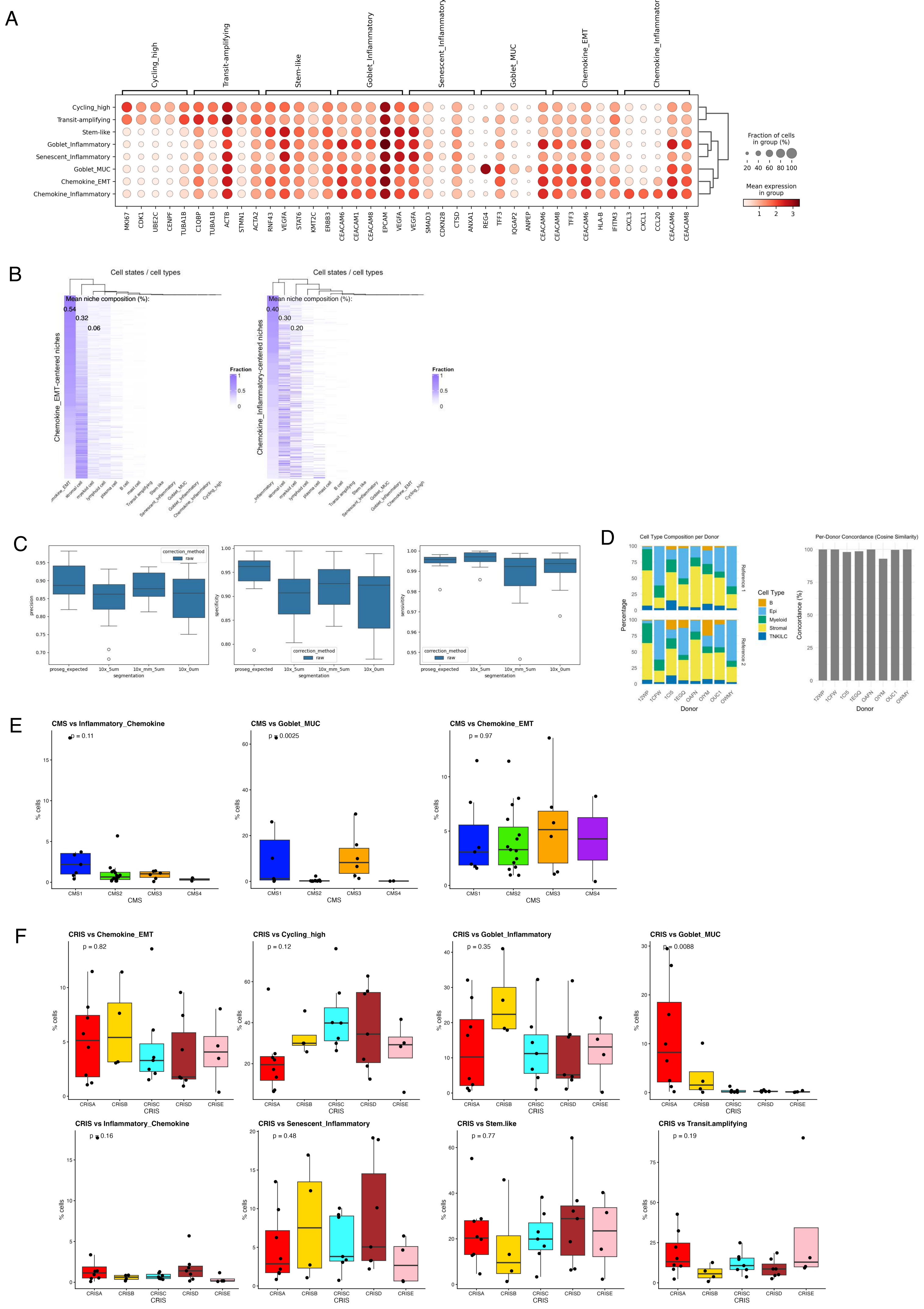

Supplementary Figure 3

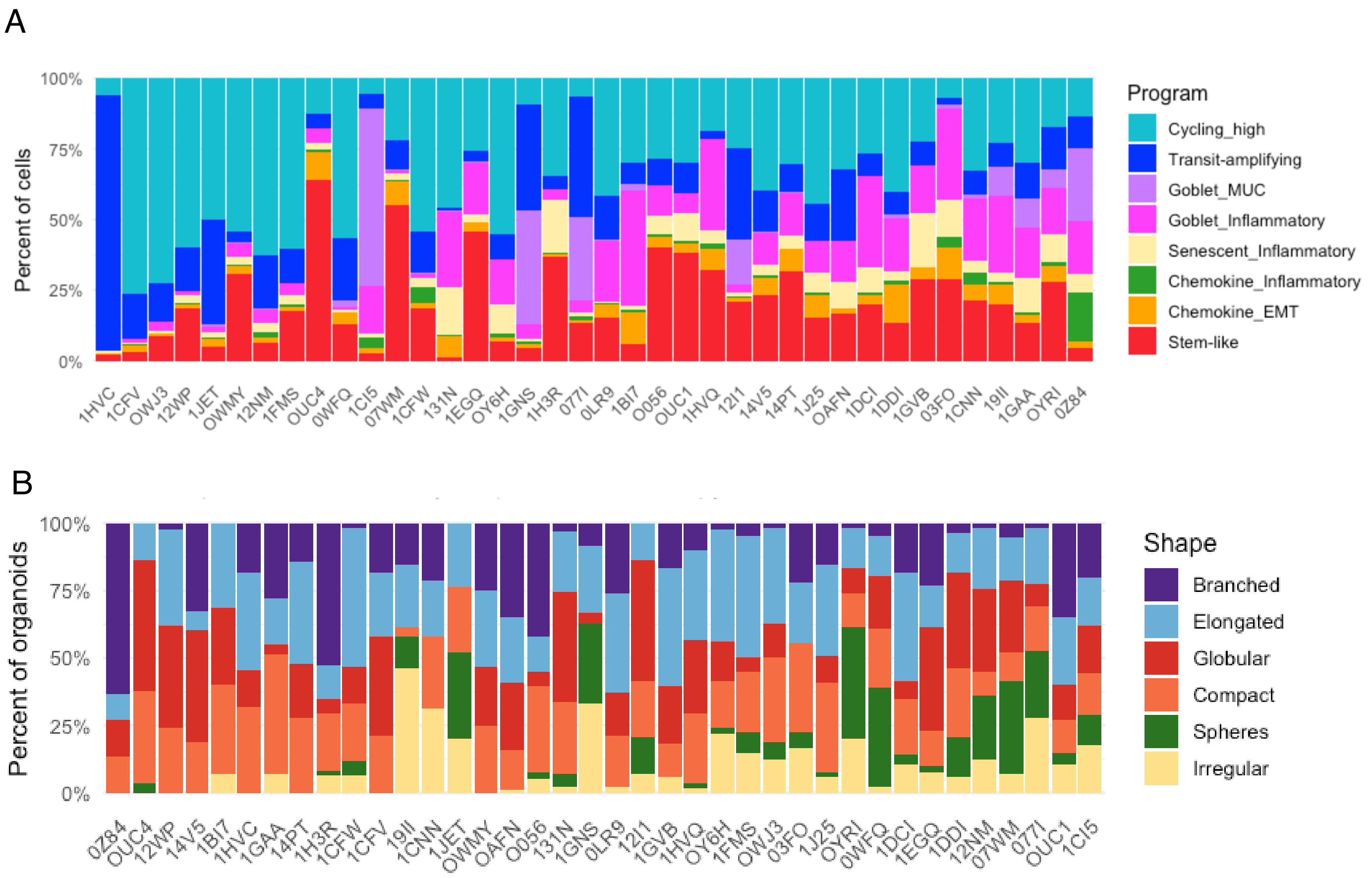

### Supplementary Figure 4

A

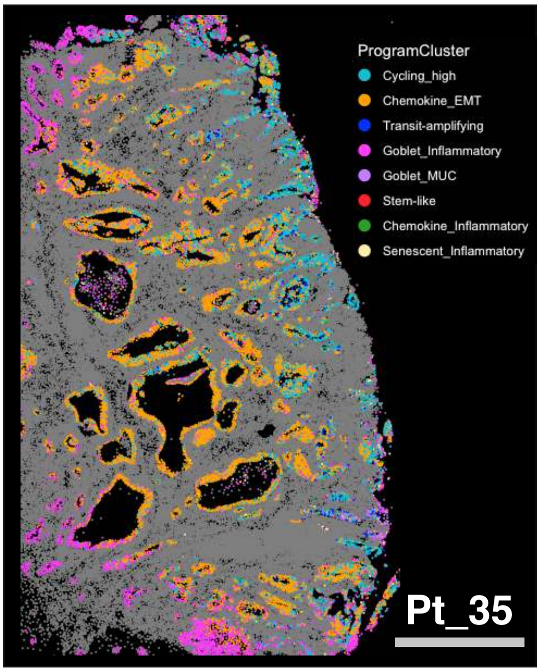


B

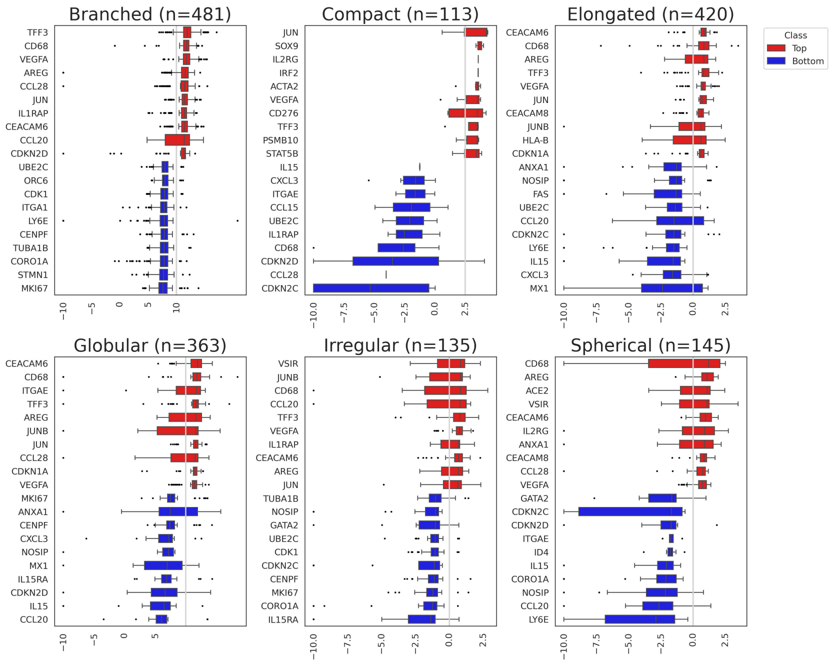


C

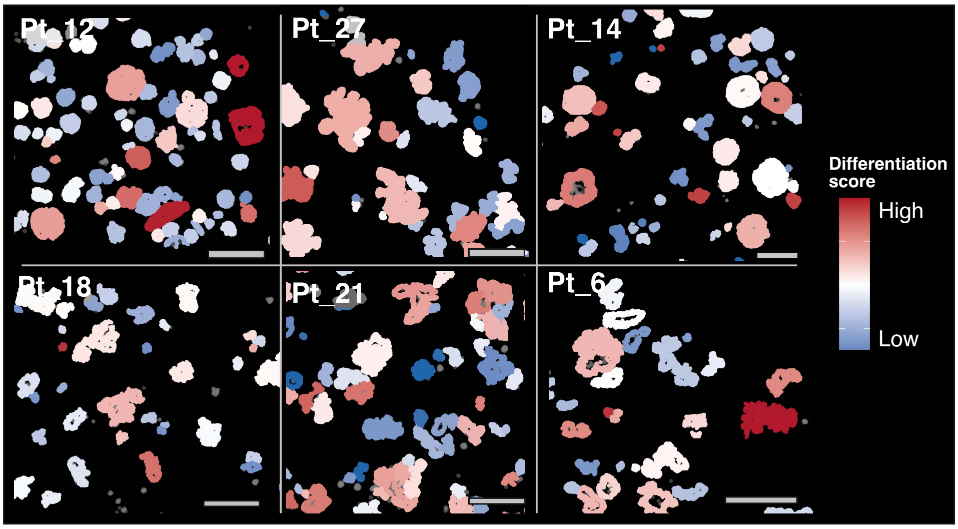

### Supplementary Figure 5

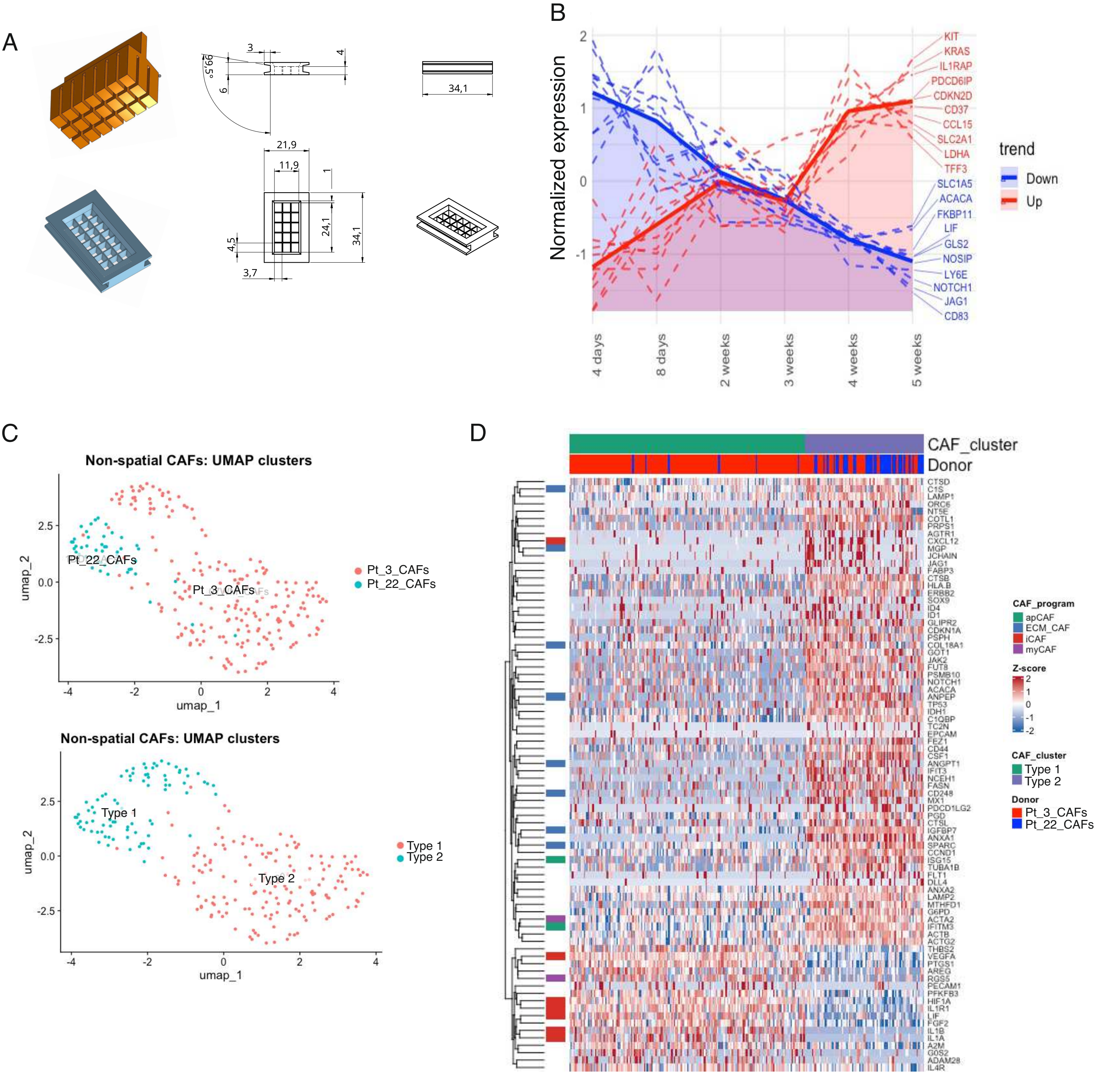
